# Single-cell transcriptome identified *ddx43^+^* cell types critical for maintenance of transient slow-cycling stem cells in planaria

**DOI:** 10.1101/2025.08.22.671706

**Authors:** Nikhil Kumar Jaligam, Mohamed Mohamed Haroon, Mainak Basu, Atriya Mazumdar, Swathi Pavithran, Vinay Kumar Dubey, Praveen Kumar Vemula, Ankit Arora, Dasaradhi Palakodeti

**Affiliations:** Institute for Stem Cell Science and Regenerative Medicine, Bangalore, Karnataka, India; Regional Centre for Biotechnology, Faridabad, Haryana, India; Cincinnati children’s hospital medical centre (CCHMC), Cincinnati, OH 45229, USA; Duke University School of Medicine, Durham, North Carolina 27710, USA

## Abstract

Cell cycle dynamics are fundamental to stem cell maintenance and differentiation. While the G1 phase is known to influence stem cell fate decisions, the mechanisms regulating its duration remain poorly understood. Using *Schmidtea mediterranea*, a highly tractable model for studying regeneration, we uncovered how the interplay of systemic and local cues from specialized cell types regulates G1 phase progression, thereby priming the cells in a slow-cycling state poised to differentiate. Single-cell transcriptomics identified G1-enriched, non-committed neoblasts that give rise to a previously uncharacterized *ddx43*⁺ cell type, localized near sub-epidermal muscle and closely associated with surrounding neoblasts. Functional analyses revealed that *ddx43⁺* cells act as gatekeepers, maintaining adjacent neoblasts in an extended G1 phase and a differentiation-primed state. *ddx43* knockdown led to neoblast hyperproliferation, highlighting their niche-like regulatory role. Following injury, extracellular signal-regulated kinase (Erk) signalling that propagates to distal regions via subepidermal muscle cells is critical for suppression of *ddx43⁺* cells by downregulating Notch signalling. This establishes a critical cross-talk between sub-epidermal muscle and *ddx43⁺* cells, mediated by the Erk–Notch axis, essential for maintaining G1 phase extension. In summary, we reveal a *ddx43*-dependent, non–cell autonomous mechanism that regulates a transient, slow-cycling neoblast population, offering new insights into how extrinsic cues orchestrate stem cell cycling during tissue homeostasis and regeneration.

## Introduction

Planarians are known for their remarkable ability to regenerate damaged or lost tissues, a capability driven by a unique population of adult pluripotent stem cells known as neoblasts (Baguñà et al., 1989; Reddien, 2018; Wagner et al., 2012). The maintenance and differentiation of specific lineages are regulated both spatially and temporally by intrinsic and extrinsic cues(Miyares & Lee, 2019; Palani & Sarkar, 2009; Paratore et al., 2002). While a significant body of work focused on understanding intrinsic factors(Petzold & Gentleman, 2021; Steens & Klein, 2022) the roles of extrinsic factors, comprising surrounding tissues and their cell types, extracellular components, and systemic cues, remain poorly understood. Previously published work has identified muscle and extracellular matrix (ECM) as key components that provide extrinsic cues critical for the maintenance and differentiation of neoblasts to specific lineages (Chan et al., 2021; Dubey et al., 2022; Gattazzo et al., 2014).

A salient feature of planarian neoblasts is their heterogeneity, primarily due to the commitment of distinct subpopulations of neoblasts to specific lineages(Adler & Sánchez Alvarado, 2015; Molina & Cebrià, 2021). Additionally, this heterogeneity is influenced by the variation in the duration of the cell cycle phases among the neoblasts. A study by Molinaro et al (2021) identified a subset of slow-cycling neoblasts characterised by their ability to retain BrdU over extended periods (Molinaro et al., 2021; Wagner et al., 2011). However, the mechanisms that maintain these slow-cycling cells in a G1/G0 phase remain unclear. In this study, we identify a mechanism that maintains a small fraction of *smedwi-1*^+^ cells in an extended G1 phase. A state maintained by their physical proximity to *ddx43^+^*cells, a population of differentiated cells associated with subepidermal muscles.

Our previous work, based on the mitochondrial content and membrane potential, identified a distinct population of neoblasts termed X2 MTG^low^ HFSC cells residing in the G1 phase. These cells exhibit enhanced pluripotency in irradiated planarian as demonstrated through rescue experiments via transplantation (Hayashi et al., 2006; Mohamed Haroon et al., 2021). Herein, we performed single-cell RNA sequencing (scRNA-seq) of X2 MTG populations followed by extensive analysis which identified a population of differentiated cells marked by the expression of *ddx43,* derived from the *smedwi-1^+^* and *ddx43^+^*neoblast.

Our study identified a small fraction of neoblasts closely associated with *ddx43^+^* cells located near the musculature. BrdU labelling experiments indicate these neoblasts reside in the extended G1 phase and are resistant to sublethal doses of irradiation. We termed this population as “transient slow cycling neoblast”. Knockdown of *ddx4*3 showed increased proliferation of these neoblasts and rendered them sensitive to sublethal doses of radiation. Together, our results suggest that this small pool of transient slow-cycling neoblasts serves as a reserve population that can re-enter the cell cycle in response to physiological stress, such as radiation-induced depletion of neoblasts.

Upon injury, the wound signal propagates to the distal tissues via longitudinal muscles through Erk signalling (Fan et al., 2023). This signalling is essential for the distal proliferation of neoblasts critical for regeneration. We found Erk signalling is required to downregulate *ddx43^+^* cells, thereby allowing the neoblast to re-enter the cell cycle. Moreover, our results also suggest that the Notch signalling downregulation is also essential for reducing the *ddx43^+^* cell numbers, further facilitating neoblast proliferation. Overall, our study highlights an essential Erk-Notch axis governs the cross-talk between muscle and *ddx43^+^* cells and is crucial for the maintenance of slow cycling neoblasts.

## Results

### Single cell transcriptome analysis identified specific cluster within X2 MTG^low^ HFSC enriched for neoblast markers

Cell populations in *Schmidtea mediterranea* (planaria) are broadly categorized into 3 classes-X1, X2 and Xins-based on their nuclear-to-cytoplasmic ratios and radiation sensitivity. The X1 population consist of actively dividing neoblasts in the S/G2/M phase of the cell cycle. The X2 population comprises neoblasts in the G1 phase and early progenitor cells (Hayashi et al., 2006). In our previous work, we employed MitoTracker^TM^-based staining to identify four subpopulations within X2 compartment based on their mitochondrial content/potential and cell size. These are classified as X2 MTG^low^ HFSC, X2 MTG^low^ LFSC, X2 MTG^high^ HFSC and X2 MTG^high^ LFSC. Through transplantation experiments into lethally irradiated planarians, we demonstrated X2 MTG^low^ HFSC represent neoblast population with enhanced pluripotency compared to other X2-derived cell types (Mohamed Haroon et al., 2021). To uncover the molecular markers enriched in X2 MTG^low^ HFSC, we performed scRNA-seq on all four X2 subpopulations **(Figure 1A)**. From this analysis, we obtained 19,015 high-quality cells, which were visualized as 16 distinct clusters using Uniform Manifold Approximation and Projection (UMAP) **(Figure 1B)** with an average of 4,500 cells per sample **(Figure 1C)**.

**Figure 1:**
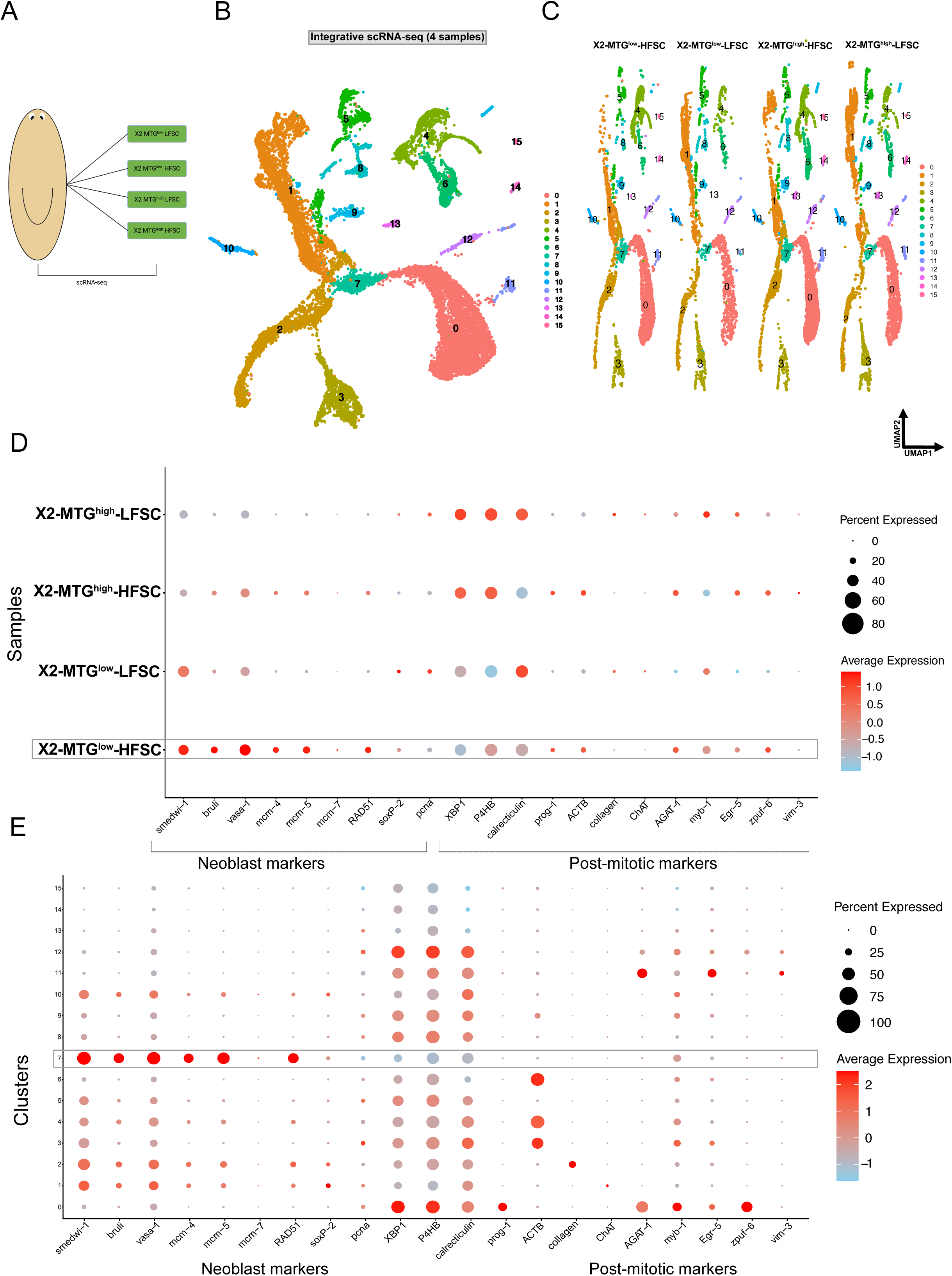
scRNA-seq illustration and expression of known markers in X2 MTG population. (A) Schematic overview of X2 MTG population from planaria (B) Uniform manifold approximation and projection (UMAP) plot represents cell clusters (n=19,015 single cells) identified from integrative analysis. (C) UMAP plot depicting cell clusters with respect to each sample namely X2 MTG^low^ HFSC, X2 MTG^low^ LFSC, X2 MTG^high^ HFSC and X2 MTG^high^ LFSC. (D) Dot plot expression representation of neoblast markers and post mitotic markers across each sample. Colors constitute average log2 expression levels scaled to unique molecular identifier (UMI) values for each cell. In it, red defines the higher expression and sky-blue defines the lower expression. Dot size represents proportionality to percentage of cells expressing particular gene. (E) Dot plot demonstrating expression of neoblast markers and post mitotic markers obtained across each clusters. Color code identify expression levels, while size of dot represents percent of cells.

We next analyzed the expression of canonical neoblast and progenitor markers across four cell populations and within individual clusters. Neoblast markers such as *smedwi-1*, *bruli*, *vasa-1*, *mcm-5* & *rad51* were markedly enriched in the X2 MTG^low^ HFSC population. In contrast, pan-differentiation markers such as *xbp1*, *p4hb* (Raz et al., 2021) were predominantly enriched in the X2 MTG^high^ HFSC & LFSC population. Dot plot analysis further confirmed that a majority of the cells in X2 MTG^low^ HFSC population exhibited high expression of neoblast markers **(Figure 1D)**. Cluster-wise analysis identified Cluster-7 as a key subset exhibiting strong expression of neoblast markers (s*medwi-1*, *bruli*, *vasa-1*, *mcm-5* & *rad51*), while lacking expression of post-mitotic or lineage committed markers. In contrast, *xbp1*, *p4hb* and other post-mitotic progenitor markers are enriched in the clusters 0 and 12 **(Figure 1E) (Figure S1A)**. As visualized using UMAP in addition to cluster 7, *smedwi-1* expression was also detected in 1,2,& 10 **(Figure 1E) (Figure S1A).**

Further, we examined the expression of known X1/X2-enriched genes across four cell populations and clusters. Markers such as *H2A, Lamin-B2, unidentified-6, DNA-helicase, vasa-2, zfp-1, Fhl-1, ap-2, khd-1* & *Imo-1* were specifically enriched in X2 MTG^low^ HFSC population and in cluster 7 **(Figure S1B & C)**. These cells lacked expression of committed progenitor marker such as *pou2/3, six1/2-2, soxP-5, nkx2.2, neuroD-1, sp6-9, runt-1, myoD* and several others **(Figure S1B & C)**. In summary, our single-cell transcriptomic analysis identifies cluster 7 as a unique sub-population within X2 MTG^low^ HFSC enriched for neoblast markers and largely devoid of lineage committed gene expression. This supports the notion that cluster 7 represents a pluripotent neoblast subpopulation with high regenerative potential.

### Gene Regulatory Network analysis (GRN) identified translation regulators and ribosome biogenesis as top candidates enriched in neoblast clusters

Single cell transcriptomic analysis identifies *smedwi-1* expression in sub-populations corresponding to clusters 1,2, 7 and 10, with highest expression levels (log2FC > 1 & adjusted p-value < 0.05) observed in clusters 1 and 7 **(Figure 2A)**. To gain insights into the molecular signature of neoblast clusters 1 and 7, we performed Gene Regulatory Network (GRN) analysis. By applying high-dimensional Weighted Gene Co-expression Network Analysis (WGCNA) (Morabito et al., 2023), we identified six modules of co-expressed genes across 4,500 cells that exhibited the highest *smedwi-1* expression **(Figure 2B)**.

**Figure 2:**
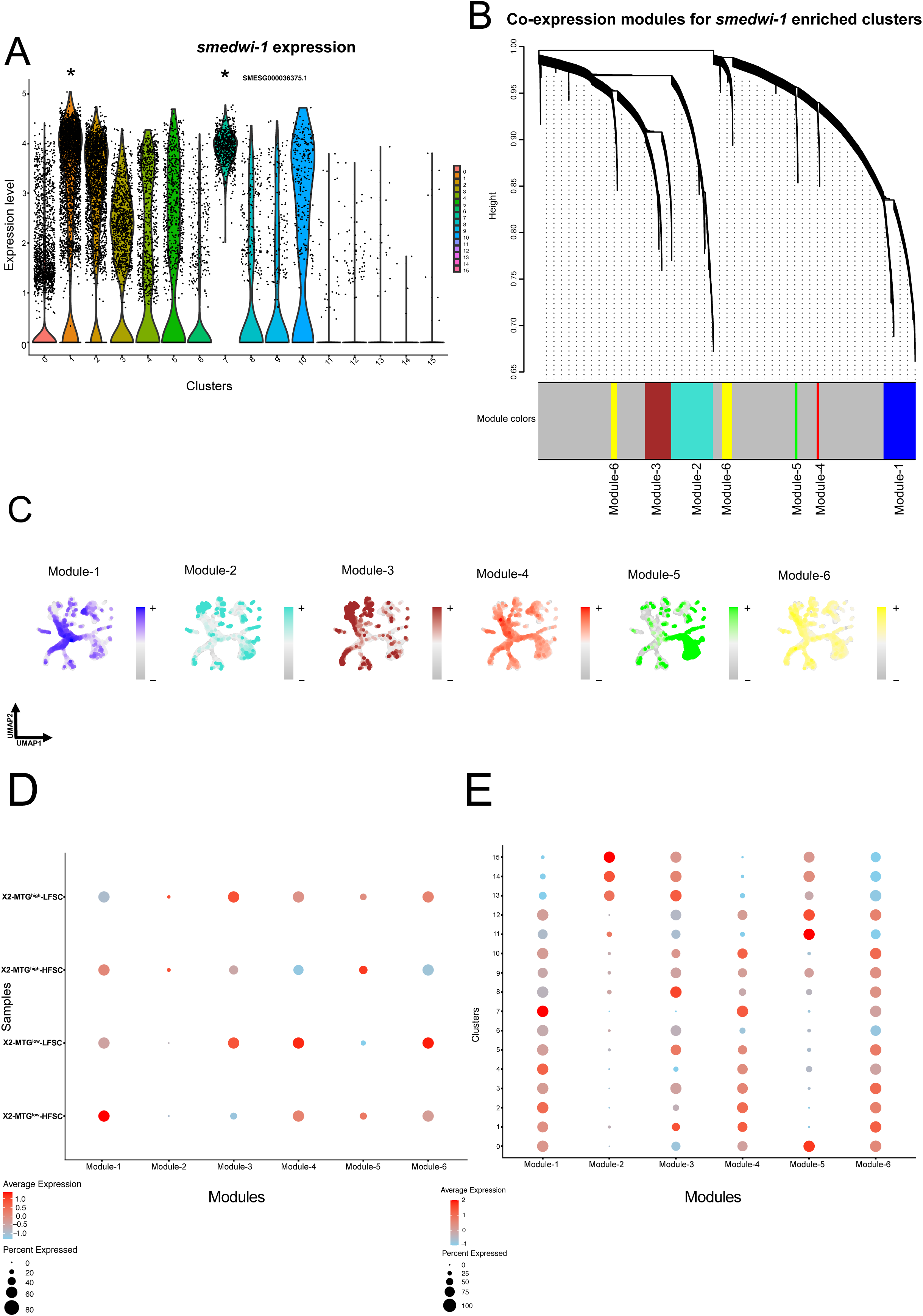
Gene Regulatory Network (GRN) analysis across neoblast clusters. (A) Volcano plot illustrating the *smedwi-1* expression across obtained cell clusters. In it star signify the significant expression of *smedwi-1* compare to different clusters (B) Weighted gene co-expression network (WGCNA) dendrogram contemplate hierarchical clustering of genes based on their expression pattern. Each leaf in dendrogram represents each gene, while the colour specified at the bottom signifies the co-expression modules. (C) UMAP plot identifying the expression pattern of different modules. Color coding explain the pattern of each module as blue for module-1,cyan for module-2, maroon for module-3,red for module-4, green for module-5 and yellow for module-6. (D) Dot-plot demonstrating average expression pattern of each module specific hub genes across each sample. In similar fashion color identify the expression levels and dot size represents the percent of cells. (E) Dot-plot representing average expression pattern of each module specific hub genes across each cluster obtained from total cells.

To visualize the distribution of these 4500 cells, we generated UMAP plots in which each module is colour coded as blue (module-1), cyan (module-2), maroon (module-3), red (module-4), green (module-5) and yellow (module-6) **(Figure 2C)**. Examining the expression pattern of genes from these six modules across the four subpopulations of X2 MTG and 16 clusters revealed that module-1 genes were most enriched in X2 MTG^low^ HFSC and cluster 7 **(Figure 2D & 2E)**. In addition, gene enrichment analysis (GO annotation) of module-1 genes identifies translation regulators as top enriched GO terms **(Figure S2A)**. This indicates that the translation regulatory mechanisms play a pivotal role in maintaining neoblast identity and function. Whereas, GO annotation from modules 2, 3, 4, 5 and 6 genes identified cilium organization **(Figure S2B)**, synapse organization **(Figure S2C)**, mRNA splicing **(Figure S2D)**, transport of ER to golgi vesicle **(Figure S2E)** and mRNA processing **(Figure S2F)** respectively. Together, our findings suggest that the molecular signatures represented in module-1 might be essential for neoblast maintenance.

### Differential expression analysis identify genes specific to pluripotent neoblast in the G1 phase of the cell cycle

In the previous section, we identified molecular signatures specific to neoblast clusters and found many of these are highly enriched in the X2 MTG^low^ HFSC subpopulation. This provides a mechanistic explanation for why the X2 subpopulation with low mitochondrial content is highly pluripotent. To further characterize this population, we performed differential gene expression analysis to identify genes specifically enriched in the X2 MTG^low^ HFSC compared to other subpopulations. Differential expression analysis identified 209 genes that were significantly enriched (Adjusted p-value less than 0.05 and log2FC greater than 1) in the X2 MTG^low^ HFSC population **(Table S1)**. Gene Ontology (GO) enrichment analysis of 209 genes revealed top enriced GO terms with molecular functions such as ribosome biogenesis, piRNA processing, Tricarboxylic Acid (TCA), metabolic process and stress granule assembly **(Figure 3A)**. We further investigated the enrichment of obtained 209 genes in 16 clusters derived from single-cell transcriptomic analysis of the X2 MTG population. Most of these genes showed enrichment in cluster 7. It also suggests that enriched genes are crucial for maintaining neoblast function **(Figure S3.1A, Figure S3.1B, Figure S3.1C, Figure S3.3A and Figure S3.3B)**.

**Figure 3:**
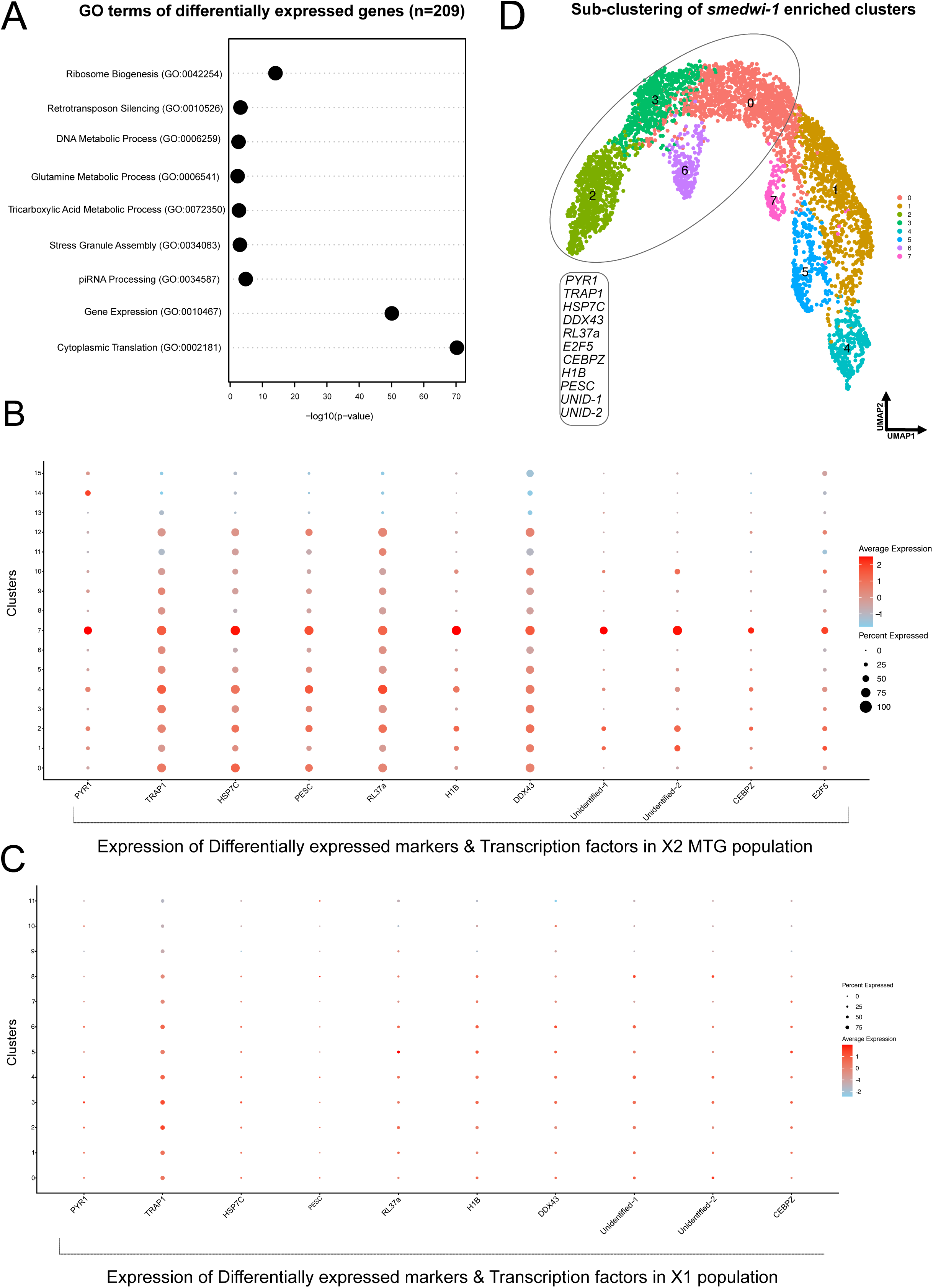
Differential expression and sub-clustering of neoblast enriched clusters. (A) Bar plot identifying the top 10 gene ontology (GO) terms obtained using differentially expressed genes (B) Dot plot depicting the expression of differentially expressed markers and Transcription factors across X2 MTG population (C) Dot plot demonstrating the expression of differentially expressed markers and Transcription factors across X1 population (D) UMAP plot describing the sub-clustering of cells from smedwi-1 enriched clusters in form of new trajectory

A subpopulation of neoblasts actively undergoing division is represented by the X1 population, characterized by high nuclear content and sparse cytoplasm, corresponding to cells in S/G2/M phase of the cell cycle. Single-cell transcriptomic analysis of X1 population identified distinct neoblast clusters, including both pluripotent and lineage committed cells (Zeng et al., 2018). Further, Raz et al. 2021 have shown that neoblasts attain differentiation marks as they progress through the S phase and undergo asymmetric division to generate progenitor cells and non-committed neoblasts in the G1 phase. To demonstrate that X2 MTG^low^ HFSC are non-committed G1 enriched neoblasts, we first compared the X1 enriched markers to the single cell transcriptome data of X2 MTG^low^ HFSC population and found that few markers, *zfp*-1, *sycp*-1, *rhombotin*-1 seem to be enriched in the neoblast cluster of X2 MTG^low^ HFSC **(Figure S3.5B)**.

Conversely, to identify genes that are specifically enriched in X2 MTG^low^ HFSC but not in X1, we examined the expression of the X2 MTG^low^ HFSC enriched genes (209 genes) across X1 neoblast clusters **(Figure S3.2A, Figure S3.2B, Figure S3.2C, Figure S3.4A and Figure S3.4B)**. Among the 209 genes, 11 genes were found to be highly expressed in X2 MTG^low^ HFSC **(Figure 3B)** and either sparsely or not expressed in X1 neoblast clusters **(Figure 3C)**. This include *pyr1, trap1, hsp7c, pesc, rl37a, h1b, ddx43, e2f5, cebpz, and* two with unidentified function *(unidentified-1, unidentified-2).* Two of the genes *cepbz* and *e2f5* are transcription factors involved in the regulation of the cell cycle and cellular differentiation. The remaining genes play a role in pyrimidine biosynthesis (*pyr1*), piRNA biogenesis and translation regulation (ATP-dependent RNA helicase (*ddx43*)), ribosome assembly and biogenesis (*rl37a*), histone packaging (*h1b*) and stress response/protein folding (*hsp7c*). However, the function of most of these genes involved in cell cycle regulation and stem cell function remains largely unknown.

Although all 11 genes were enriched in cluster 7–*pyr1, h1b, unidentified-1, unidentified-2, cepbz and e2f5* – were exclusive to cluster 7, while others (*trap1, hsp7c, pesc, rl37a, ddx43)* were also expressed to lesser extent in the clusters marked by post mitotic markers **(Figure 3B)**. Furthermore, sub-clustering based on *smedwi-1* expression identified eight clusters with *ddx43, trap1, pyr1,rl37a and* e2f5 enriched in clusters 0,2,3 & 6 **(Figure 3D) (Figure S3.5A)**. This confirms that the 11 genes are part of the G1 enriched neoblast.

We next performed pseudotime analysis to investigate the dynamic trajectories of these cell clusters. From our analysis, we identified cluster 7 as a prominent cluster expressing neoblast markers, while lacking expression of pan-differentiation markers (*xbp1, p4hb*) (Raz et al., 2021). Using ‘Monocle’ (Qiu et al., 2017), we set cluster 7 as the root to define lineage trajectories. The directionality of pseudotime from high to lower pseudo-time was validated using *smedwi-1,* which showed decreasing expression along the axis. We then used additional markers including *xbp1, p4hb*, *prog-1 and vasa-1* to track their expression pattern along the pseudotime. Neoblast markers followed the trend of higher to lower pseudo time, whereas the differentiation markers followed the opposite trajectory **(Figure 4A)**. We obtained the list of Fate Specific Transcription Factors (FSTF) from H. O. King et al., (2024) & Raz et al., (2021) and checked the expression along the pseudo-temporal pattern. The epidermal markers such as *p53* and *soxP-3* followed the expression pattern of lower to higher pseudotime, whereas the markers for muscle such as *myoD* followed the opposite pattern from higher to lower pseudotime **(Figure 4B)**.

**Figure 4:**
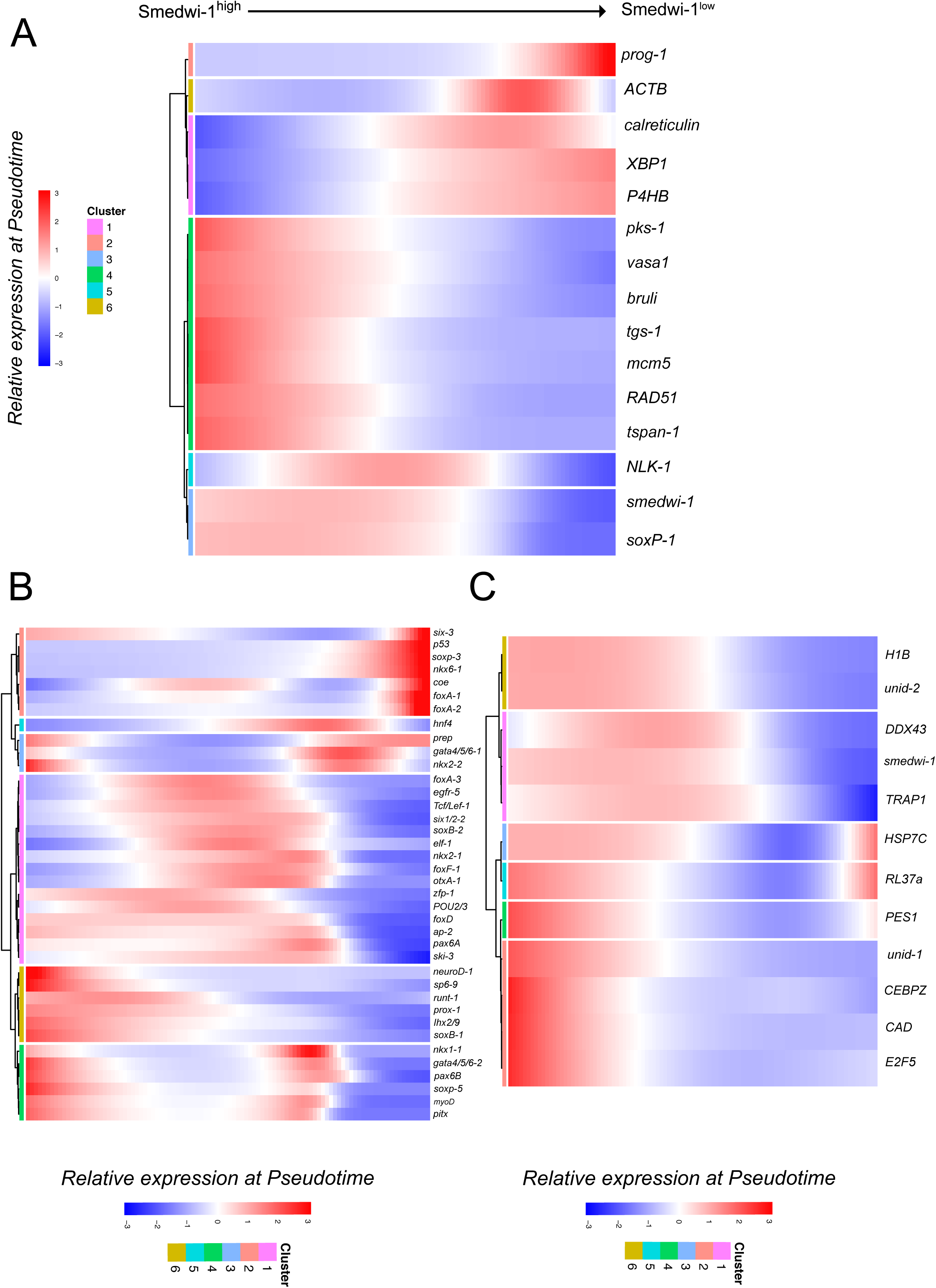
Neoblast and Post mitotic markers modulates the pseudotime trajectory inference. (A) Heatmap showing the dynamics of markers changing gradually over *smedwi-1* higher expression to lower expression. Genes (row) and cells (column) are arranged according to the pseudotime progression. Genes mentioned in first set are represented for the post mitotic markers and second set of genes are represented for neoblast markers. (B) Heatmap exhibiting pseudotime progression of Fate Specific Transcription Factors (FSTF) obtained from (H. O. King et al., 2024; Raz et al., 2021). Therein epidermal markers showed pseudotime expression from lower to higher whereas muscle, intestinal markers showed opposite trend from higher to lower expression (C) Heatmap depicting the trajectory inference of 11 genes enriched in X2 MTG population in compare to X1 population. Likewise all the eleven genes follow the similar pseudotime progression like neoblast marker as *smedwi-1*.

Further, we checked pseudotime progression of 11 genes enriched in X2 MTG^low^ HFSC, which followed pseudo-temporal progression similar to *smedwi-1*, transitioning from higher to lower pseudotime **(Figure 4C)**. To infer lineage trajectories, we performed Slingshot analysis (Street et al., 2018), using cluster 7 as the root. This revealed a bifurcating trajectory from cluster 7, with one branch representing a neoblast lineage and the other a post-mitotic lineage, as visualized in the UMAP **plot (Figures S4A–S4C)**.

In summary, our in-depth single-cell transcriptomic analysis of X2 MTG population identified 11 genes specifically enriched in X2 neoblast, representing the G1 phase of the cell cycle. While these genes appear to be important for neoblast maintenance, their precise role in regulation of G1 phase and pluripotency remains to be elucidated.

### RNAi Screening identify unique role of *ddx43* and *h1b* in negative regulation of neoblast proliferation in planaria

To investigate the role of 11 genes enriched in the X2 MTG^low^ HFSC population, we performed RNA interference (RNAi) mediated knockdowns. Animals were fed with dsRNA at a final concentration of 250 ng/µL mixed with beef liver extract. A total of six feeds were administered at three-day intervals, followed by post-treatment observations for 7 days **(Figure 5A)**. Five out of 11 knockdowns resulted in lethal phenotypes at varying time points. Among the 11 genes, *e2f5*, *pesc*, *unidentified 2 (un2)*, and *cebpz* knockdowns caused lesions leading to animal lysis, while knockdown of *rl37a* resulted in body paralysis, followed by lysis **(Figure S5A)**. Animals that did not exhibit phenotypic defects were fixed seven days after the final feed for further analysis. Quantitative PCR was performed to validate the RNAi efficiency **(Figure S5B).**

**Figure 5:**
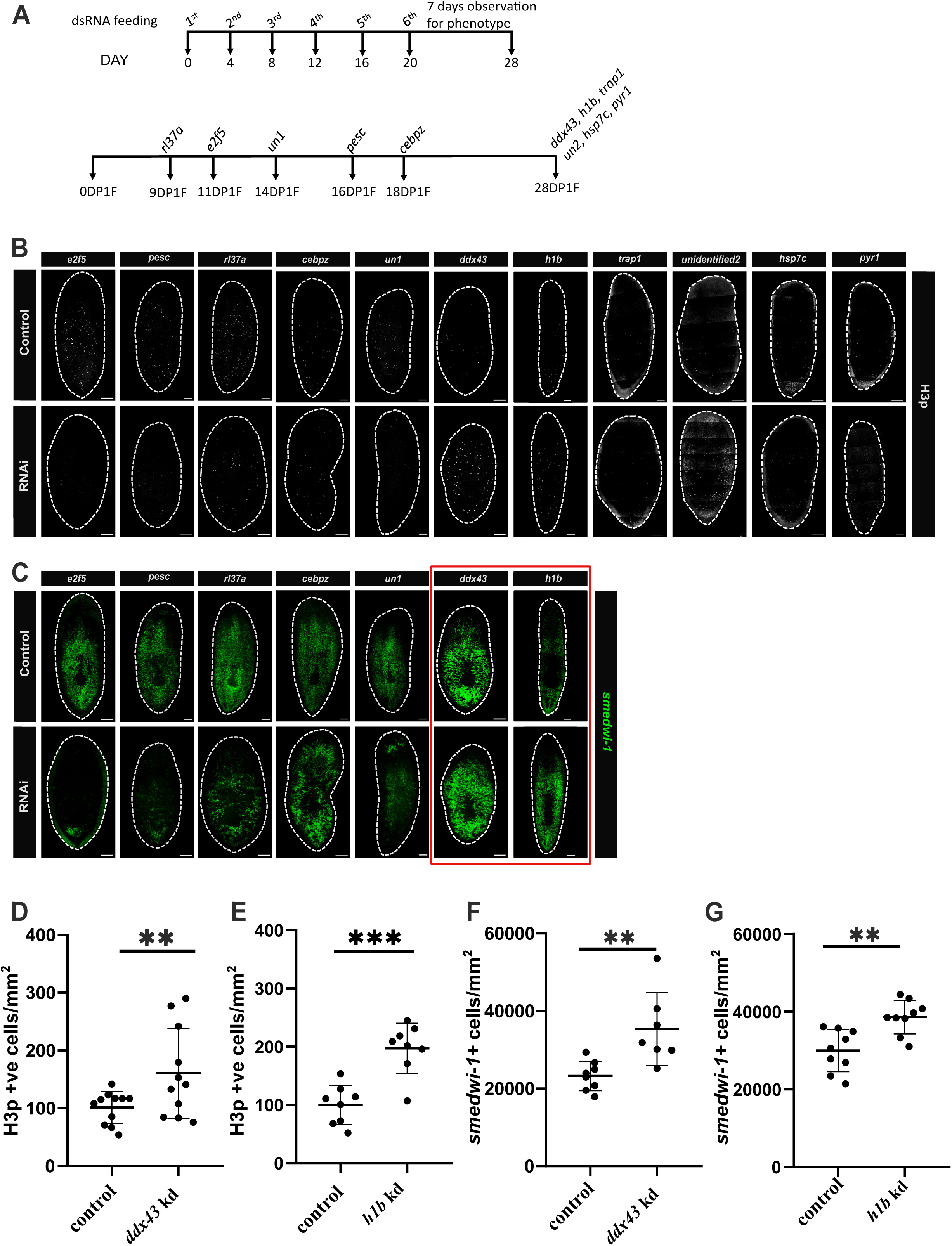
RNAi screening revealed hyperproliferation in *ddx43* and *h1b* knockdown animals. (A) Schematic representation of the RNAi knockdown strategy applied to screen 11 candidate genes. Animals were fed with gene-specific dsRNA six times with 3-day interval between feeding. Animals exhibiting phenotypes were fixed immediately upon observation, while those without observable phenotypes were fixed at 28^th^ day. (B) Immunostaining using anti-phospho-histone H3 (H3p) for all eleven gene knockdowns. Scale bars:100 µm. (C) RNA-FISH for *smedwi-1* staining in gene knockdowns of *e2f5*, *pesc*, *rl37a*, *unidentified-1 (un1), ddx43*, and *h1b*. Scale bars: 100 µm. (D, F) Quantification of H3p^+^ and *smedwi-1*^+^ cells in *ddx43* knockdown animals. p value: ** <0.01; *** <0.001 (E, G) Quantification of H3p^+^ and *smedwi-1*^+^ cells in *h1b* knockdown animals. p value: ** <0.01

Immunostaining for phosphorylated histone H3 (H3P), a mitotic marker, revealed a reduction in proliferating cells in *e2f5*, *pesc*, *rl37a*, *cebpz*, and *unidentified 1 (un1)* knockdowns **(Figure 5B, S5C)**. In contrast *ddx43*, *h1b*, *trap1,* and *un1* knockdowns showed an increase in H3P^+^ cells indicating altered proliferative dynamics **(Figure 5B, 5D, 5E and S5C)**.To further assess the impact on neoblast populations, we performed RNA fluorescence in situ hybridisation (RNA-FISH) for *smedwi-1*, a neoblast marker **(Figure 5C)**. *e2f5*, *pesc*, *cebpz*, and *un1* knockdowns resulted in a reduction of neoblasts, whereas *rl37a* knockdown led to neoblast clustering or compacting. Although *trap1* and *un2* knockdowns showed an increase in mitotic cells **(Figure S5C)**, while neoblast numbers remained unchanged **(Figure S5D)**. Interestingly, *ddx43* and *h1b* knockdowns exhibited a significant increase in the number of *smedwi-1^+^* cells (**Figure 5F, 5G)**. Together, our results show increased numbers of H3P^+^ and *smedwi-1^+^*cells, suggesting hyperproliferation of neoblasts in *ddx43* and *h1b* knockdown animals **(Figure S5E)**.

### *ddx43*-dependent regulation of *h1b* establishes hierarchical relationship critical in controlling neoblast proliferation

Given that both *ddx43* and *h1b* knockdowns result in a similar hyperproliferation phenotype, a relatively uncommon outcome in neoblast-targeted RNAi screens suggesting a role in restraining stem cell proliferation. We investigated whether these genes are expressed within the same cell types, which might explain the similar phenotype between both gene knockdowns. Analysis of our single-cell transcriptomic data revealed a negative correlation between *ddx43* and *h1b* expression, indicating limited co-expression between *ddx43* and *h1b* **(Figure S6A)**. To better understand this relationship and its phenotypic consequences, we examined their spatial expression patterns in planarians by performing a series of double fluorescence in situ hybridisation (dFISH) experiments using probes against *ddx43* or *h1b* along with neoblast marker *smedwi-1* in wild-type (WT) animals. RNA-FISH for *ddx43* and *smedwi-1* revealed *ddx43* is expressed in *smedwi-1^+^* neoblasts and differentiated cells. These differentiated cells expressing *ddx43* are in physical proximity and are surrounded by neoblasts **(Figure 6A)**. Similarly, dFISH for *h1b* and *smedwi-1* revealed ∼50% of *h1b^+^* cells co-express *smedwi-1* **(Figure 6B)**. We next explored lineage relationship of *ddx43*⁺ and *h1b⁺* cell types using scRNA-seq from X2 MTG population. UMAP analysis revealed 92.58% of *smedwi-1⁺* cells in the X2 MTG population express *ddx43*. Among these, a subset (55.11%) co-expresses *h1b*, forming a distinct *smedwi-1⁺/ddx43⁺/h1b⁺* triple-positive population **(Figure S6B)**. Additionally, we identified a separate *smedwi-1⁺/h1b⁺* population (∼55%) that lacked *ddx43* expression. These findings suggest a hierarchical lineage structure, wherein *smedwi-1⁺/ddx43⁺* neoblasts represent a parent lineage that gives rise to both *smedwi-1⁺/ddx43⁺/h1b⁺* and *smedwi-1⁺/h1b*⁺ progeny **(Figure S6B)**. This lineage relationship was further validated by dFISH experiments **(Figure 6C)**. We observed co-expression of *smedwi-1* with both *ddx43* and *h1b*, as well as co-expression of *ddx43 and h1b* in a subset of cells. These findings suggest *ddx43^+^ and h1b^+^* cell types might have originated from the same parent lineage. Further, using dFISH experiments for *ddx43* and *h1b* reveals *h1b^+^* cells are in close physical proximity to *ddx43*⁺ cells **(Figure 6C)**. This observation prompted us to test whether *ddx43*^+^ cells might regulate *h1b^+^* cells in a non-cell autonomous or hierarchical manner. To test this, we analyzed *h1b* expression in *ddx43* knockdown animals and vice versa **(Figure 6D, 6E)**. Strikingly, the number of *h1b*⁺ cells were significantly reduced in *ddx43* knockdown, while *ddx43^+^* cells remained unaffected in *h1b* knockdown **(Figure 6F, 6G)**. This suggests that *ddx43* acts upstream of *h1b* and plays a critical role in supporting the *h1b^+^*cell population. These results are further corroborated by QRT-PCR analysis, wherein *h1b* transcript levels were reduced by 50% in *ddx43* knockdown animals **(Figure S6C)**. In contrast, *ddx43* transcript levels remained unchanged in *h1b* knockdown animals **(Figure S6D)**. Further, bulk transcriptome sequencing of *ddx43* knockdown animals followed by bioinformatics analysis revealed increased expression of differentiated markers in the knockdown animals compared to the controls **(Figure S6E)**. This suggests that *ddx43^+^* cell types might play a pivotal role in maintaining the adjacent neoblast in extended G1 phase, which were primed to differentiate upon entering the cell cycle.

**Figure 6:**
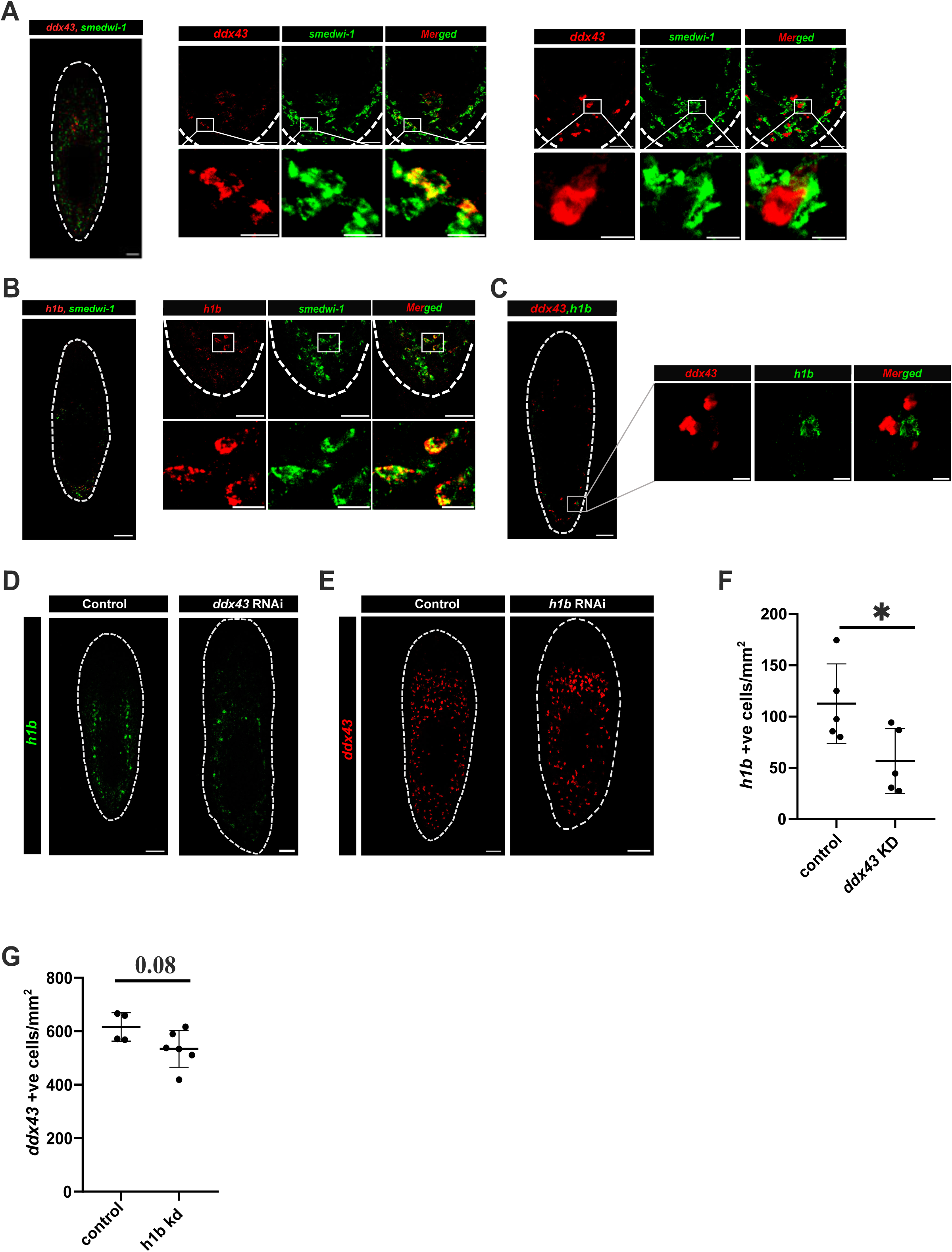
In-situ staining conferred *ddx43* is essential for *h1b* expression. (A) RNA-FISH signiting by using *ddx43* and *smedwi-1* probes in wild-type animals. First insert shows co-localization of *ddx43* with *smedwi-1* in the tail region. Second inset shows *smedwi-1*^+^ cells surrounding *ddx43*^+^ cells in the tail region. (B) RNA-FISH staining by using *h1b* and *smedwi-1* probes in wild-type animals. Inset shows *h1b* colocalizing with *smedwi-1* in the tail region. (C) RNA-FISH staining using *ddx43* and *h1b* probes in wild-type animals. Inset shows *ddx43*^+^ cells in close proximity to *h1b*^+^ cells. (D) RNA-FISH for *h1b* in *ddx43* knockdown animals. (E) RNA-FISH for *ddx43* in *h1b* knockdown animals. (F, G) Quantification of *h1b*^+^ and *ddx43*^+^ cells in *ddx43* and *h1b* knockdowns. Scale bars: Whole-animal: 100 µm; tail insets: 50 µm; single cell insets: 10 µm. p value: * <0.05

Together, our data reveal a hierarchical regulatory relationship between *ddx43* and *h1b*, where *ddx43* promotes *h1b* expression and function. This axis appears to be essential for controlling neoblast proliferation under homeostasis condition.

### *ddx43* positive cells play pivotal role in maintaining nearby neoblasts in slow-cycling state and confer resistance to sublethal doses of radiation

To investigate the cell cycle state of the neoblasts in close physical proximity to *ddx43*⁺ cells, we performed RNA-FISH using probe against *ddx43 and* immunostaining using antibody against H3p, a marker of cells in the G2/M phase of the cell cycle. Only a small fraction (∼10%) of proliferating neoblasts located near the *ddx43^+^* cells are H3p^+^ which further suggests that these neoblast are predominantly in the G1 phase and may be slow cycling.

In an effort to directly access whether these neoblasts are indeed in the slow-cycling state, we conducted a long-term BrdU pulse chase experiment. Planarians were fed with BrdU and examined for label retention in neoblast on 15 days post feeding (dpf). We performed immunofluorescent staining with anti-BrdU antibody followed by dFISH using probes against *smedwi-1* and *ddx43*. Strikingly, neoblast in the close proximity to *ddx43^+^*cells retained BrdU signal, indicating a slow-cycling nature **(Figure 7A)**.

**Figure 7:**
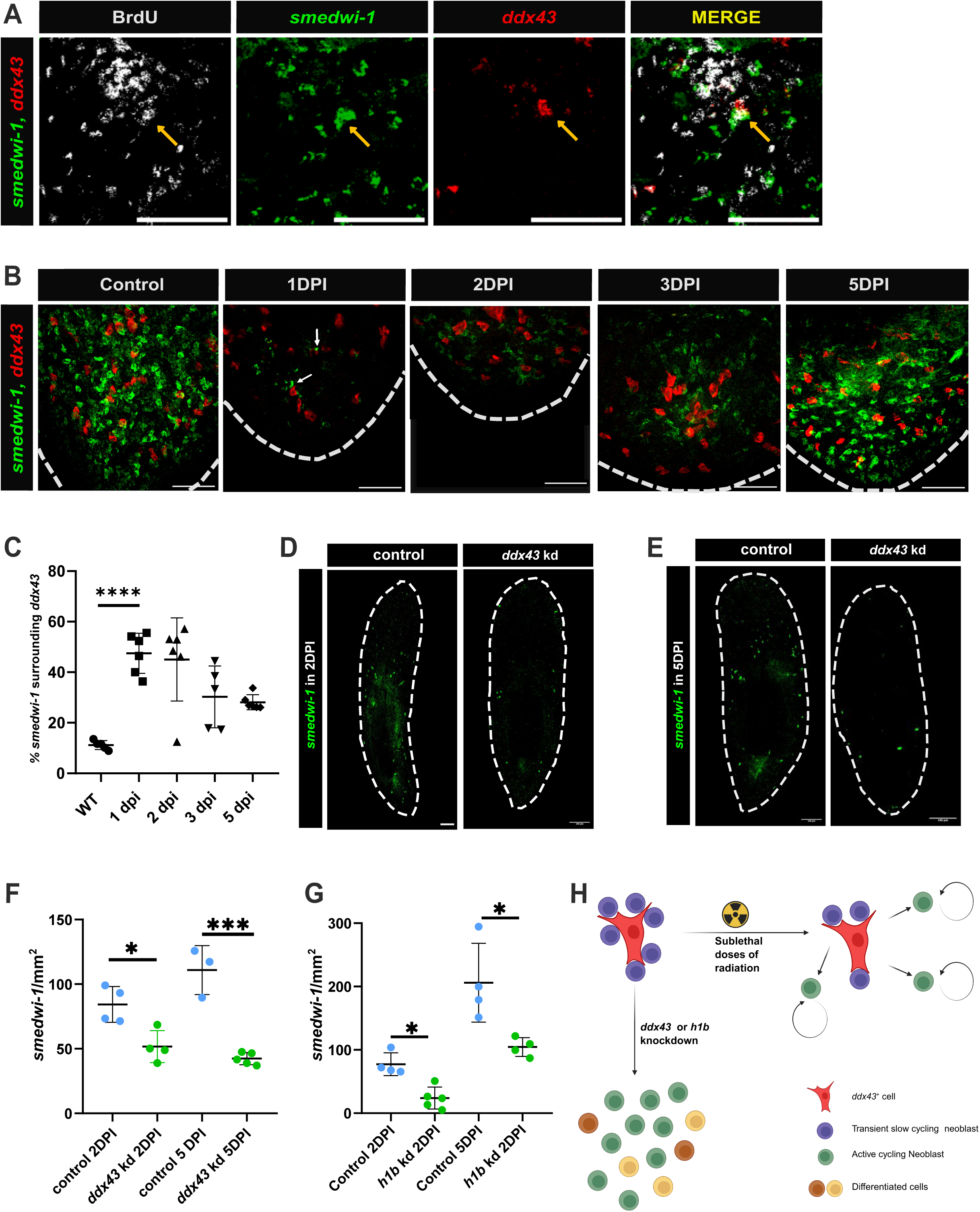
Sublethal doses of irradiation exhibit *ddx43* importanat for transient slow cycling neoblasts. (A) Triple labelling of *ddx43*, *smedwi-1* (RNA-FISH), and BrdU immunostaining in 15-day BrdU chased animals. Arrow indicates *smedwi-1*^+^BrdU^+^ cell near a *ddx43*^+^ cell. (B) RNA-FISH for *ddx43* and *smedwi-1* in wild-type and sub-lethally irradiated animals (15 Gy) at 1, 2, 3, and 5 days post-irradiation (DPI). Gamma radiation was used to irradiate the animals. (C) Quantification illustrating *smedwi-1*^+^ cells in proximity to *ddx43*^+^ cells at indicated timepoints as 1, 2, 3 and 5 dpi along with wildtype animals after irradiation.*smedwi-1^+^*cells in physical proximity to *ddx43^+^* cells are considered and are divided with total *smedwi-1^+^* cells and multiplied with 100 to obtain the percentage. p value: **** <0.0001 (D, E) RNA-FISH for *smedwi-1* at 2dpi and 5dpi in *ddx43* knockdown animals subjected to sublethal radiation (15Gy). (F, G) Quantification of *smedwi-1*^+^ cells at 2DPI and 5DPI in *ddx43* and *h1b* knockdowns. Each condition having four animals. p value: *<0.05; ***<0.001 (H) Model depicting *ddx43*^+^ cells maintaining transient, slow-cycling neoblasts. Loss of *ddx43* or *h1b* leads to hyperproliferation. While upon sublethal doses of irradiation, slow-cycling neoblasts re-enter the cell cycle and proliferate.

When planarians were exposed to sublethal dose of radiation (15 Gy) **(Figure S6F)**, partial loss of neoblasts was observed. Existing literature suggests that slow cycling cells are resistant to radiation (Lyle & Moore, 2011; Skvortsova et al., 2015). This led to our hypothesis that neoblasts resistant to sublethal doses of radiation constitute a slow cycling population residing near the *ddx43^+^* cells. To test this, we irradiated the planarians with 15 Gy and performed dFISH using probes against *smedwi-1 and ddx43.* At 1-day post-irradiation (dpi), the surviving neoblasts were predominantly located near *ddx43*⁺ cells. Notably, by 3 dpi, many of the neoblasts had migrated away from *ddx43*⁺ cells, likely to repopulate the tissue **(Figure 7B, 7C)**. These findings suggest that neoblasts surviving sublethal doses of radiation are slow-cycling in nature which are maintained in the vicinity of *ddx43^+^* cells.

To determine whether *ddx43*⁺ cells are essential for maintaining the radiation resistant and slow-cycling neoblast population, we performed RNAi-mediated knockdown of *ddx43* followed by sublethal doses of radiation and stained with *smedwi-1* **(Figure 7D, 7E)**. At 2 and 5 dpi, *ddx43* knockdown animals exhibited a significant reduction in surviving neoblasts, indicating compromised radiation resistance **(Figure 7F, 7G)**. These results demonstrate that *ddx43*⁺ cells are critical for the survivability of neoblast to sublethal doses of radiation by maintaining them in transient slow-cycling state. In summary, our findings highlight a pivotal role of *ddx43*⁺ cells in regulating the local microenvironment to maintain neighbouring neoblast in a transient slow-cycling state **(Figure 7H)**. However, the molecular mechanisms by *which ddx43*⁺ cells sustain this state and how this regulation is modulated in response to injury remains to be elucidated.

### Erk signalling downregulate *ddx43*^+^ cells by suppressing Notch, critical for neoblast proliferation

A recent study by Fan et al., (2023) demonstrated Erk signalling propagates rapidly from the site of injury towards the distal end along the longitudinal muscle. This signalling was shown to be essential for neoblast proliferation occurring within 6 hours post amputation (hpa), in the distal from the site of injury. We know that *ddx43^+^* cells negatively regulate proliferation and ERrk promotes proliferation, therefore, we hypothesised that Erk signalling might deplete *ddx43^+^* cells to promote proliferation in distal tissues. To investigate the fate of *ddx43*⁺ cells and their associated neoblast following injury, we amputated the anterior region of the planaria just below the head and fixed them at 6 hpa. We performed FISH using probes against *ddx43* **(Figure 8A)**. In the proximal region near the wound, *ddx43*⁺ cell numbers remained unchanged comparable to homeostasis **(Figure 8B)**. In contrast, *ddx43^+^* cell numbers were significantly reduced in the distal region **(Figure 8C)**. These findings led us to hypothesize that the Erk-dependent neoblast proliferation at the distal end, as reported in Fan et al. (2023) might be facilitated by the depletion of *ddx43^+^* cells. This in turn could enable the nearby neoblast to exit slow-cycling state and enter proliferation. To test this, we treated regenerating animals with the Erk inhibitor FR180204 (25 µM conc.), which blocks the activity of phosphorylated Erk. dFISH with *ddx43* and *smedwi-1 probes* revealed that Erk-inhibition significantly increased *ddx43*⁺ cell numbers in the distal region compared to untreated controls **(Figure 8D, 8E)**. The *ddx43*^+^ cell numbers in Erk inhibitor condition remain unchanged in proximal regions compared to untreated control **(Figure S7A)**. These results indicate that Erk signalling is required to reduce *ddx43*⁺ cell numbers, thereby promoting neoblast proliferation.

**Figure 8:**
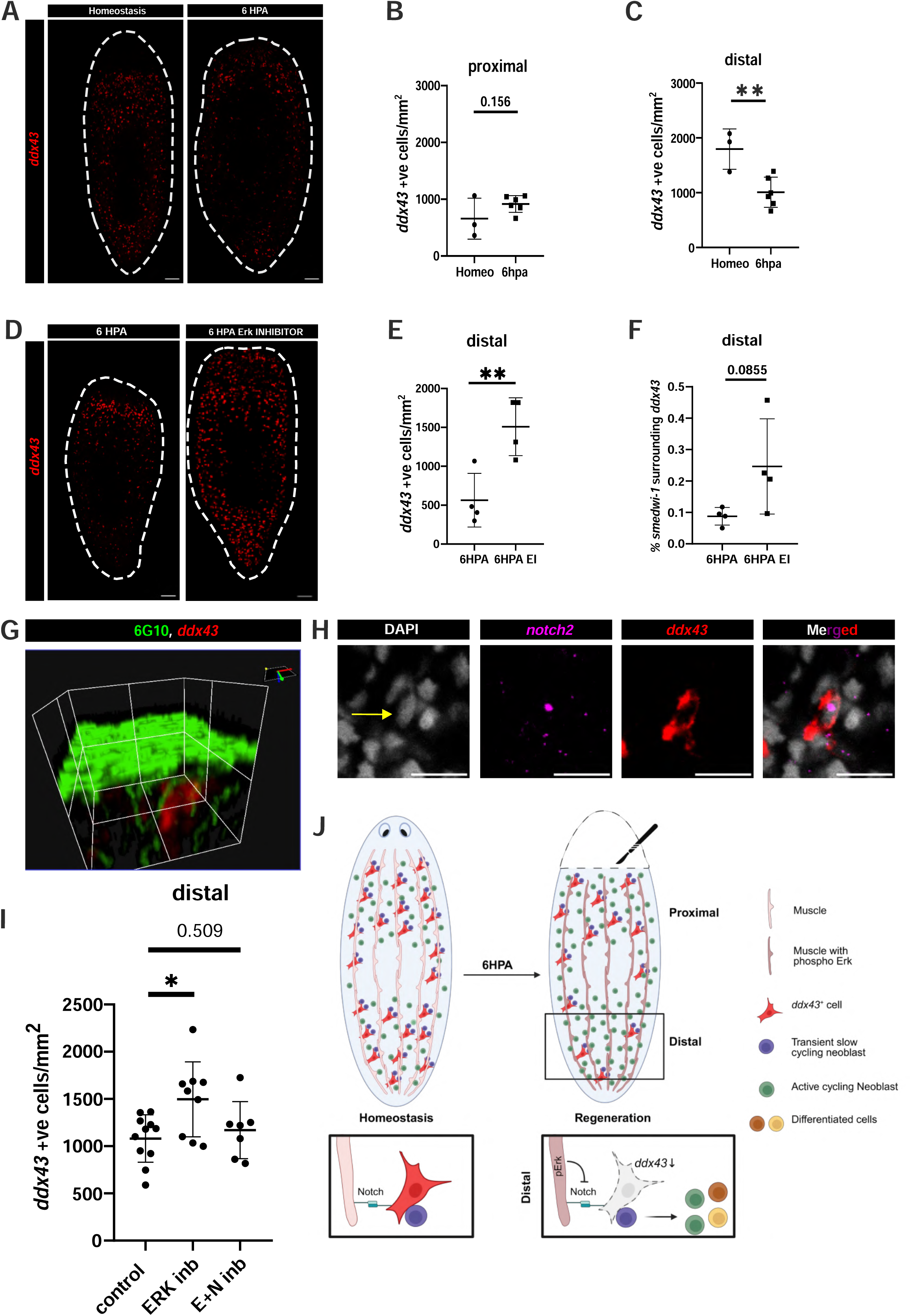
Erk activity crucial for downregulation of *ddx43* in distal region. (A) RNA-FISH for *ddx43* in homeostasis and 6 hours post-amputation (6 hpa). Heads were amputated just below the eyes and fixed after 6 hours. (B, C) Quantification of *ddx43*^+^ cells in proximal and distal regions in uncut and 6 hpa condition. In homeostasis conditions, the prepharynx is compared with proximal region of 6HPA animal and postpharynx region is compared to distal region of 6hpa animal. p value: **<0.01 (D) RNA-FISH showing for *ddx43* at 6 hpa in untreated and Erk inhibitor (FR180204) treated animals. (E) Quantification of *ddx43*^+^ cells in distal regions of planaria in untreated and Erk inhibitor treated animals. p value: ** <0.01 (F) Quantification showing *smedwi-1*^+^ cells in close proximity to *ddx43+* cells in distal regions of untreated and Erk-inhibited animals. *smedwi-1^+^*cells in physical proximity to *ddx43^+^* cells are considered and are divided with total *smedwi-1^+^* cells and multiplied with 100 to obtain the percentage. Each condition exhibiting four animals. (G) 3D image rendering of muscle (6G10) and *ddx43* stain showing the close association of muscle with *ddx43^+^* cells. The 3D redering was done in FV3000 software from Olympus. (H) RNA-FISH for *ddx43* and *notch2* in wild-type animals. *notch2* expression was observed in *ddx43^+^* cells. Scale bars: 10 µm. (I) Quantification of *ddx43*^+^ cells in the distal region at 6 HPA in untreated, Erk-inhibited, and Erk+Notch (E+N) co-inhibited animals. (J) Model illustration for *ddx43+* cells during injury. Upon injury, Erk-mediated Notch signalling leads to downregulation of *ddx43^+^* cells in distal regions post-injury, and promotes proliferation.

Next, we quantified the proportion of neoblasts near the *ddx43*⁺ cells in both control and Erk-inhibited conditions. A significantly higher fraction of neoblasts were found in the proximity to *ddx43*⁺ cells in Erk-inhibited animals **(Figure 8F)**. Given that Erk signalling in planaria is known to act via muscle cells (Fan et al., 2023), we investigated spatial relationship between *ddx43*⁺ cells and muscle. Co-staining for the muscle markers, 6G10 and *ddx43* in uninjured wild-type animals reveals that *ddx43*⁺ cells are in close proximity with the muscle cells **(Figure 8G, S7B, S7C)**. Together, these results suggest that Erk signalling plays a pivotal role in downregulating the *ddx43^+^* cell number in the distal region during regeneration crucial for neoblast proliferation.

To further explore the signalling cross talk between muscle and *ddx43^+^*cells, we analyzed single cell transcriptomic data to identify signalling pathway, using genes enriched in *ddx43^+^/smedwi-1^-^* cells. Notably, *notch2* is highly expressed in *ddx43^+^/smedwi-1^-^*cells suggesting Notch signalling may play a role in maintaining *ddx43^+^* cells **(Table S2)**.Previous studies have shown that Notch signalling regulate stem cell dynamics in planarians. For instance, inhibition of Notch using DAPT in *Dugesia japonica* have shown to increase neoblast proliferation by 6hpa (Dong et al., 2021)*. Furthermore,* dFISH of *notch2* with *ddx43* confirmed the expression of *notch2* in *ddx43^+^* cells **(Figure 8H, S7D, S7E)** and *notch2* KD also showed increased proliferation **(Figure S7F, S7G)**. This observation led to our hypothesis that Erk signalling may suppress *ddx43^+^* cells by inhibiting Notch activity.

To test this hypothesis, we treated animals with both Erk and Notch inhibitors and examined *ddx43^+^* cell numbers in the distal region from the site of injury. Strikingly, *ddx43^+^* cells were significantly reduced under dual Erk and Notch inhibition, reaching comparable levels comparable to untreated controls **(Figure 8I)**. In summary, our data suggests that Erk signalling, transmitted via muscle following injury suppress Notch activity in *ddx43^+^* cells, leading to a reduction in the cell numbers. This down regulation is a necessary step to facilitate the proliferation of adjacent neoblast during regeneration. In summary, the pERK signalling downregulates notch in distal tissues, thereby, depleting *ddx43^+^* cell number, an essential step for neoblast proliferation **(Figure 8J)**

## Discussion

Cell cycle regulation in stem cells has traditionally been viewed as a mechanism to expand the stem cell pool, essential for their maintenance. However, emerging evidence across diverse systems has shown that the duration of the cell cycle, particularly the G1 phase, acts as an active regulatory node in stem cell renewal and fate specification. The G1 phase acts as a critical window during which stem cells integrate extrinsic and intrinsic cues to determine whether to self-renew, enter quiescence, or undergo lineage commitment (Dalton, 2015; Pardee, 1974; Zaveri & Dhawan, 2018). In human and mouse embryonic stem cells (ESCs), dynamic changes in the G1 duration have been shown to influence fate outcomes. For instance, shortened G1 was associated with maintenance of naïve pluripotency, driven by extrinsic signals such as Leukemia Inhibitory factor LIF (Coronado et al., 2013), while prolonged G1 phase increases sensitivity to differentiation cues like retinoic acid (Pauklin & Vallier, 2013). G1-specific chromatin accessibility also appears to prime cells for fate transitions (Sela et al., 2012; Trouth et al., 2025). Furthermore, molecular regulators like Cyclin E1 and p27^Kip1^ have been shown to directly link G1 progression with stem cell maintenance and differentiation (Chetty et al., 2015; Menchón et al., 2011).

Despite these insights, the link between G1 duration and differentiation is not universally consistent. Li et al., (2012) demonstrated that mere elongation of G1 in mouse ESCs via overexpression of cell cycle inhibitors like p21 and p27 did not accelerate differentiation. Conversely, shortening G1 by overexpressing cyclins did not delay it. These findings suggest that G1 length alone may be insufficient to dictate fate decisions in certain contexts and that additional regulatory mechanisms likely buffer stem cells from differentiation cues.

In this study, we used *Schmidtea mediterranea*, a model organism with robust regenerative capabilities, to interrogate how cell cycle dynamics regulate the maintenance and function of adult pluripotent stem cells. Planarians maintain a population of adult pluripotent stem cells, a sub-population of neoblasts, capable of regenerating all somatic cell types. Our study identified a transient slow-cycling population of neoblasts with an extended G1 phase. These cells appear to exist in a state of differentiation readiness, without being lineage committed and may play a crucial role in facilitating differentiation.

Further, we found that this slow-cycling G1 neoblasts are in close proximity to previously uncharacterized *ddx43-*expressing *cell type*, which we term “Janitor cells”-gatekeepers of cell cycle. The discovery of *ddx43*⁺ cell was serendipitous arising from our search for molecular markers specific to G1-phase of the neoblast. To characterize G1 neoblast population, we focused on the X2 MTG^low^ HFSC subset described by Haroon et al., (2021), a G1-enriched neoblast population. The earlier study identified four distinct populations of X2 based on mitochondrial content and cell size: X2 MTG^low^ HFSC, X2 MTG^low^ LFSC, X2 MTG^high^ HFSC, and X2 MTG^high^ LFSC. Among these, X2 MTG^low^ HFSC cells, characterized by low mitochondrial content and larger cell size exhibited the highest pluripotency, as evidenced by their ability to rescue lethally irradiated animals upon transplantation.

Building on these insights, we isolated all four X2 MTG subpopulations and performed single-cell transcriptomic sequencing to identify G1-phase-specific signatures. Differential gene expression analysis comparing X2 MTG^low^ HFSC cells to the other X2 populations and X1 neoblasts (S/G2/M phase cells) revealed 11 genes specifically enriched in G1-phase neoblasts. Additionally, we also checked the pseudo-temporal trajectory differences, which concur with the biological trajectory of non-committed neoblasts and the post-mitotic markers. Our obtained eleven X2 enriched genes showed expression of higher to lower pseudo-time similar to neoblast pattern suggesting the co-expression of these genes in the G1 enriched neoblast population. Among these, *ddx43* exhibited distinct expression pattern of its expression in the *smedwi-1^+^* and in differentiated cells. Lineage tracing based on the single transcriptome data revealed that *ddx43^+^*cells types arise from a *ddx43^+^/smedwi-1^+^* G1 enriched neoblast population.

To investigate the functional relevance of *ddx43^+^* cell types, we performed *ddx43* knockdown, which resulted wide-spread neoblast hyperproliferation. This suggests that *ddx43*⁺ cells exert non-cell autonomous control over neoblast cycling. Remarkably, these transient slow cycling cells adjacent to *ddx43*⁺ cells are resistant to sub-lethal radiation and repopulate the animals with neoblasts, indicating to a protective niche-like function of *ddx43*⁺ cells. Together, our data highlight a critical role for spatial cues from *ddx43*⁺ cell in regulating stem cell dynamics, particularly in the G1 phase, essential for tissue homeostasis. Importantly, the transient slow-cycling neoblasts described are distinct from the slow-cycling cells reported by Molinaro et al., (2021), which were characterized by a metabolically distinct profile. In contrast, the population we described here are microenvironment-dependent, shaped by local interactions with *ddx43*⁺ cells.

To explore the role of *ddx43*⁺ cells during regeneration, we performed anterior amputations and analyzed the neoblast response at 6 hours post amputation (hpa), corresponding to the first mitotic peak (Wenemoser & Reddien, 2010). We observed a marked reduction in *ddx43*⁺ cell numbers at the distal end, but not the proximal end of the wound. This inversely correlated with the findings of Fan et al., (2023), who showed Erk-dependent proliferation of neoblast at the distal end of the wound. we hypothesized that Erk signalling may promotes neoblast proliferation by suppressing *ddx43⁺* cells. This was tested by treating the anterior amputated animals with a well-known Erk inhibitor, which led to a significant decrease of the *ddx43*⁺ cell types in the distal region of the wound. This supports our hypothesis that Erk signalling is a critical for regulation of cells marked by *ddx43*.

Fan et al., (2023) also showed that Erk activation propagates through the longitudinal muscle from the injury site towards the distal end. We speculated that Erk-mediated downregulation *ddx43⁺* cell types requires proximity to muscle tissue. Indeed, our data showed that the *ddx43*⁺ cells are closely surrounded by the subepidermal muscle, suggesting juxtacrine signalling between muscle and *ddx43*⁺ cells. To identify the signalling component within the *ddx43^+^* cells that might respond to Erk, we analysed single-cell transcriptome data from the *ddx43-*enriched cells, which identified Notch2 as a potential candidate. This also led to the hypothesis that Erk activation modulates *ddx43^+^*cell types by altering Notch function.

Previous studies have shown interactions between Erk and Notch pathways in various systems. For example in mouse cochlea, Mek/Erk signalling supresses hair cell differentiation by maintaining Notch activity, while Erk inhibition downregulates Notch components (Ma et al., 2024). In breast cancer, MAPK-ERK controls Notch activation via regulation of Jagged1 (JAG1), and Erk inhibition disrupts Notch-driven tumor progression (Izrailit et al., 2013). Similarly, in oral squamous cell carcinoma, Erk activation induces Jagged1 expression, activating the Notch pathway and enhancing cancer stemness (L. J. Li et al., 2022). These examples suggest a conserved Erk–Notch axis, which may also operate in planarians to coordinate injury responsive niche function. To test this, we performed dual inhibition of Erk and Notch signalling. This restored *ddx43*⁺ cell numbers in distal end from the site of amputation to control levels, unlike Erk inhibition alone, which led to increased levels *ddx43*⁺ cells. These results establish the Erk-Notch axis as a critical regulatory mechanism modulating *ddx43*⁺ cell function during early wound response and regeneration.

Beyond injury response, our study also reveals a unique mechanism for maintaining the neoblast in an extended G1 phase. While Raz et al., (2021) showed that planarian neoblasts can initiate fate specification during the S/G2/M phases of the cell cycle without compromising potency, our findings highlight an additional regulator layer. Here, extrinsic cues from *ddx43⁺* cells maintain neighbouring neoblasts in a poised, pluripotent and slow-cycling G1 state. These neoblasts are not fate primed but serve as a reserve population, capable of rapid activation upon injury, enhancing regenerative resilience.

In conclusion, our findings reveal that G1 phase regulation is an active, microenvironment-mediated mechanism of stem cell maintenance in planaria. The spatial association between *ddx43*⁺ cells and neoblasts adds a new dimension to our understanding of how regenerative capacity-not solely regulated by intrinsic transcriptional program, but also by extrinsic, niche-derived restraint. Rather than a passive interval, G1 extension emerges as a key regulatory feature supporting stem cell identity and regenerative potential.

## Methods

**Table 1.**
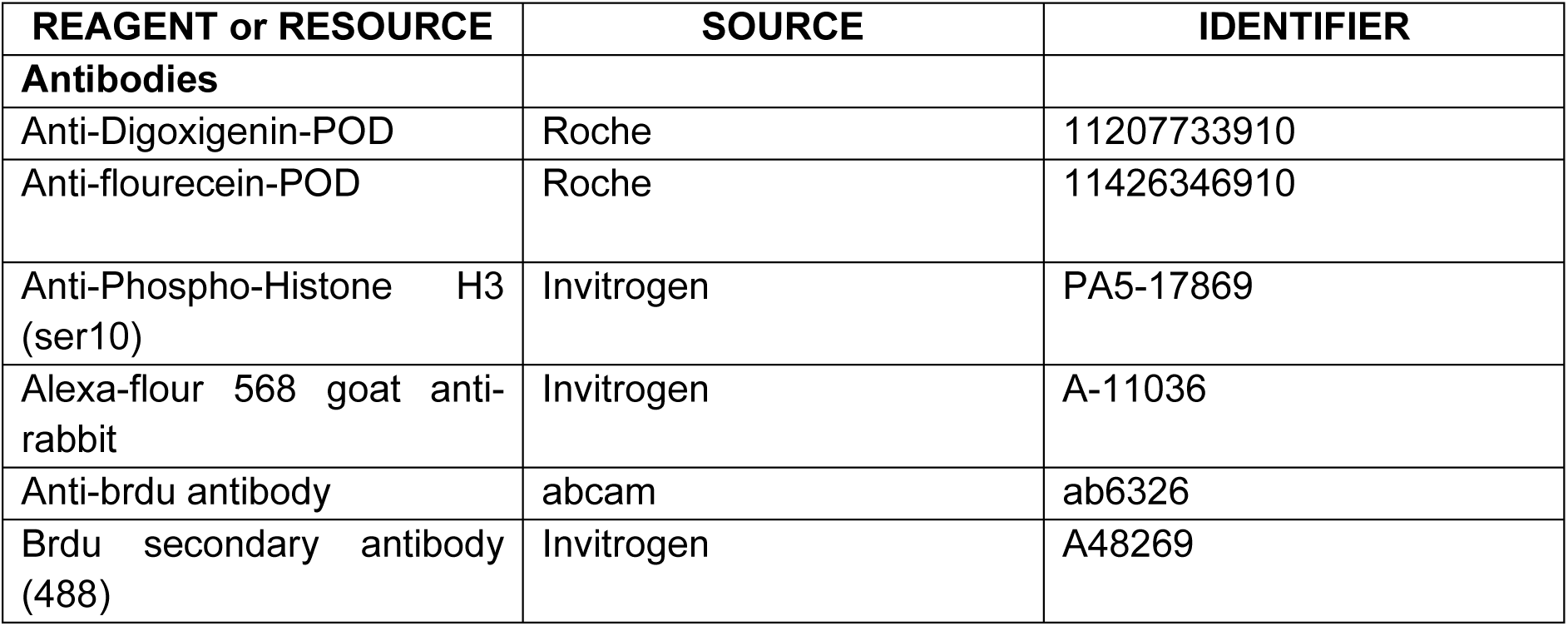

| REAGENT or RESOURCE | SOURCE | IDENTIFIER |
| --- | --- | --- |
| **Antibodies** |  |  |
| Anti-Digoxigenin-POD | Roche | 11207733910 |
| Anti-flourecein-POD | Roche | 11426346910 |
| Anti-Phospho-Histone H3 (ser10) | Invitrogen | PA5-17869 |
| Alexa-flour 568 goat anti-rabbit | Invitrogen | A-11036 |
| Anti-brdu antibody | abcam | ab6326 |
| Brdu secondary antibody (488) | Invitrogen | A48269 |

**Table 2.**

| Chemicals, (peptides and recombinant proteins | SOURCE | IDENTIFIER |
| --- | --- | --- |
| Sodium azide | Sigma-Aldrich | S2002 |
| TRizol Reagent | Thermo Fisher | 15596018 |
| Dimethyl sulfoxide (DMSO) | Sigma-Aldrich | D8418 |
| ERK Inhibitor II, FR180204 | Sigma-Aldrich | 865362-74-9 |
| DAPI | Invitrogen | D1306 |
| T7 polymerase | NEB | M0251S |
| Thermostable inorganic phosphate | NEB | M0296L |
| Recombinant RNase inhibitor | TAKARA | 2313A |
| Dithiothreitol (DTT) | INVITROGEN | Y00147 |
| 20X SSC | Invitrogen | 15557044 |
| Hydrogen peroxide | Fisher Scientific | 7722-84-1 |
| Tween 20 | Sigma-Aldrich | P9416 |
| Deionized formamide | AMBION | AM9342 |
| RNase-free DNase I | NEB | M0303L |
| Formaldehyde | Sigma-Aldrich | F8775 |
| Triton X-100 | Sigma-Aldrich | T8787 |
| Proteinase K | Genetix Biotech Asia | PG-6070 |
| Ethidium Bromide | AMRESCO | X328 |
| BRDU | MERCK | 10280879001 |
| N-acetyl-L-cysteine | SIGMA ALDRICH | A7250 |
| Kanamycin | HIMEDIA | MB105 |
| DIG RNA labelling mix | Roche | 11277073910 |
| Flourescein RNA labelling mix | Roche | 11685619910 |
| PowerUP SYBRGreen master mix | Thermo Fisher Scientific | 4367659 |

**Table 3.**

|  |  |  |
| --- | --- | --- |
| **Commercial assays** |  |  |
| Chromium Next GEM Single Cell 3' Reagent Kits v3.1 | 10X genomics | PN-1000121 |
| cDNA kit | TAKARA | 6110A |
| wizard plus sv minipreps dna purification system | Promega | A1460 |
| TA cloning kit | Invitrogen | K207040 |
| QIAprep spin Miniprep kit | QIAGEN | 27104 |
| MEGAclear Transcription Clean-Up Kit | Thermo scientific | AM1908 |
| NEBNext® Poly(A) mRNA Magnetic Isolation | NEB | E7490L |
| NEBNext® Ultra™ II Directional RNA Library Prep Kit | NEB | E7765L |

**Table 4.**

|  |  |  |
| --- | --- | --- |
| **Software and algorithms** |  |  |
| cellranger software | v6.0.0 | 10X Genomics |
| Seurat | v4.0.0 |  |
| R | v4.0.3 |  |
| Monocle | v2 |  |
| Slingshot | v2 |  |
| hdWGCNA | v0.2.19 |  |
| WGCNA | v1.72-1 |  |
| Fiji | v1.54p | https://fiji.sc/ |
| Graphpad Prism | v8.0.1 | https://www.graphpad.com/ |
| Inkscape | v1.4 | https://inkscape.org/ |
| Biorender |  |  |

**Table 5.**
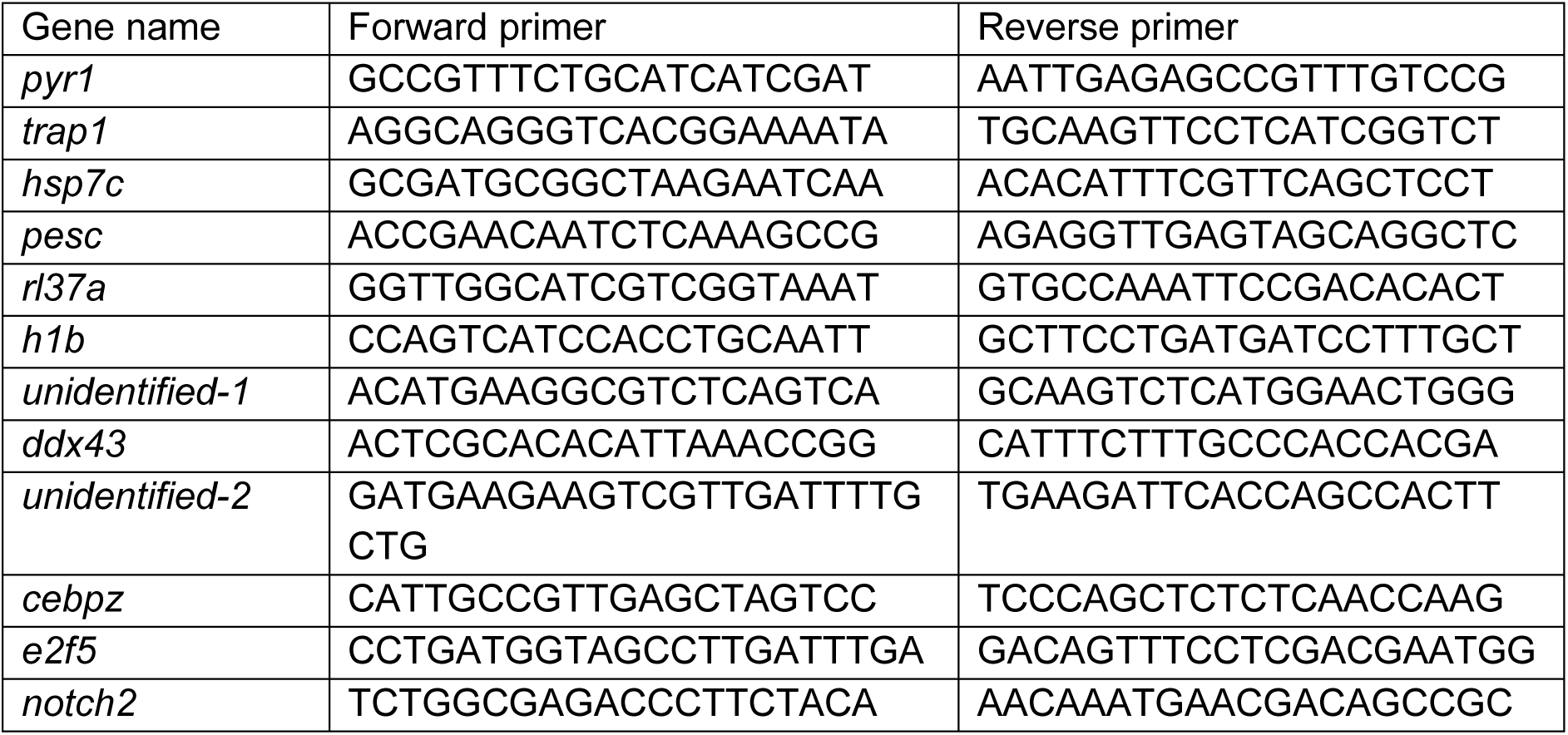
Primers used for dsRNA and Riboprobe synthesis.

**Table 6.**
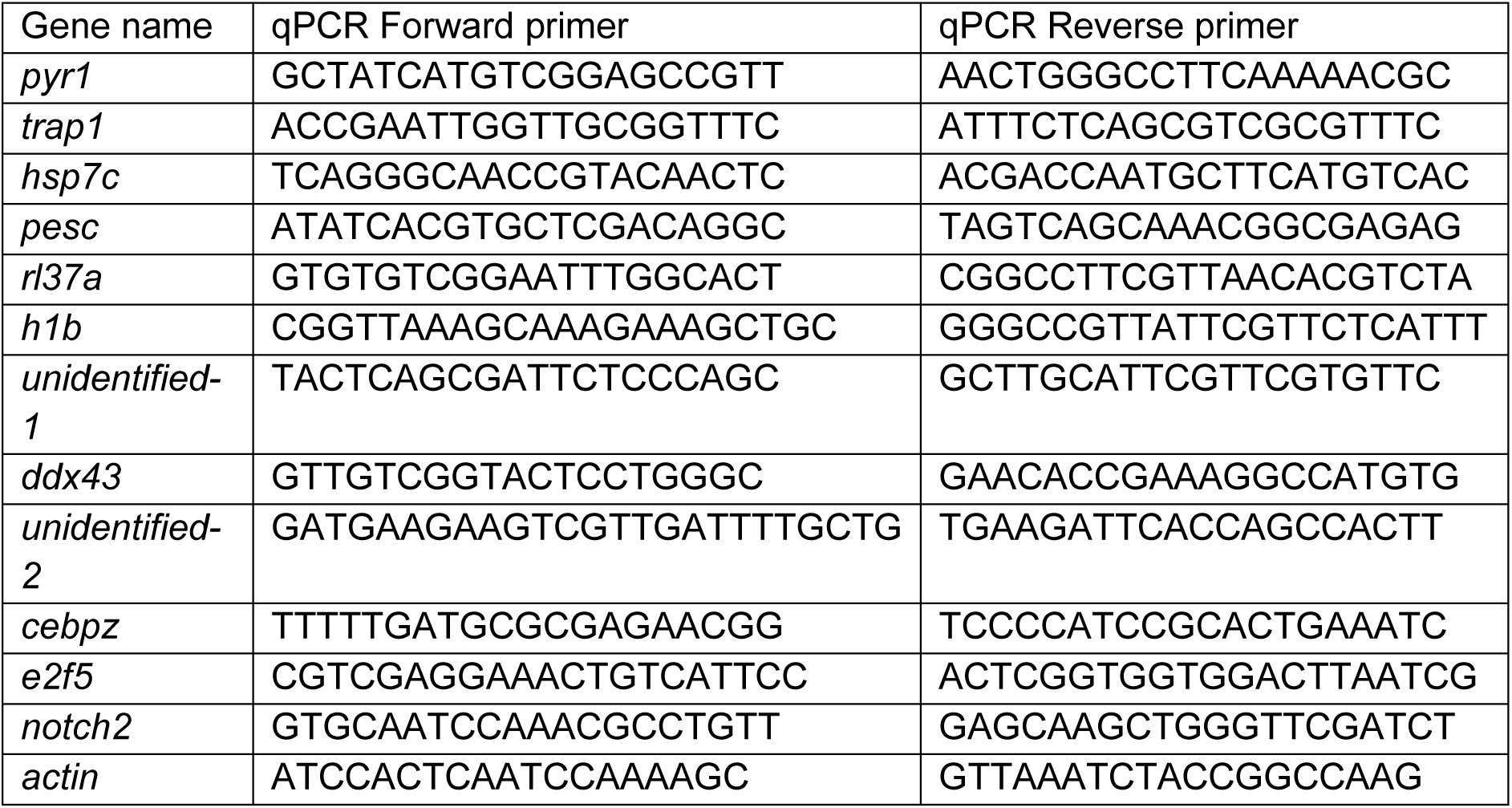
Primers used for qPCR.

### Animal husbandry

The planarian species ‘*Schmidtea mediterranea’* was used for our study. All animals were maintained in 1x Montjuïc water at 20°C in dark conditions within Sanyo incubators. ph range of 7-7.4 had been maintained for the 1x Montjuïc water. Planaria were fed with beef liver extract at every 4^th^ or 5^th^ day. The animals were starved for a week before taking them for the experiment.

### Single-cell RNA-sequencing library preparation

Staining and FACS sorting of X2 MTG populations were performed as described previously (Haroon et al., 2022; Mohamed Haroon et al., 2021). Briefly, ∼0.7 cm planaria were used and the animals were diced into small pieces using a scalpel in CMFB+1% Bovine serum albumin (BSA). After chopping, the fragments were transferred to a 50 mL centrifuge tube using a wide bore pipette tip. Cells were mechanically dissociated by repeated pipetting and the resulting suspension is filtered using 40um cell strainer. The strained solution is centrifuged at 290g for 10mins at 4 degree. The supernatant is discarded and the cells were resuspended in isotonic planaria media (IPM) with 10% Fetal Bovine Serum (FBS). The cells were then stained with 40ug/ml hoechst 33342 for 50 mins at Room Temperature (RT). Then Mito Tracker Green (MTG) was added at 100 nM concentration for 20 mins at RT. After staining, the sample is centrifuged at 290g for 10mins at 4 degree and then cells were resuspended in IPM+10% FBS. The samples were immediately analysed through flow cytometry. The X1 and X2 gates were set based on Hoechst staining. X1 MTG populations and X2 MTG were gated as described in Haroon et al., (2022) and Haroon et al., (2021). Sorting was performed in BD FACS ARIA III cell sorter, using a 100 µm nozzle with the sheath fluid pressure set at 20 psi. The sorted cells were counted using a haemocytometer, and the viability was tested using trypan blue exclusion assay. The sorted cells had more than 90% viability. The cells were then subjected to single cell RNA (scRNA) sequencing library preparation using 10X Chromium Next GEM Single Cell 3ʹ Reagent Kit v3.1 according to the manufacturer’s instruction. Sequencing was done using Ilumina Nova seq 6000.

### scRNA-seq data processing

Raw scRNA-seq reads were aligned to the planaria genome *Schmidtea mediterranea* S2F2 (SMESG.1) genome using 10X Genomics CellRanger count (v6.0.0) (Zheng et al., 2017) to attain gene/cell count matrices. The corresponding genome and General Feature Format (GFF) were obtained from planmine database (Rozanski et al., 2019) and prepared with CellRanger ‘mkref’ function. Each sample in form of UMI count were merged into a table and imported into Seurat (v4.0.0)(Butler et al., 2018) (Stuart et al., 2019) within the R environment (v4.0.3). Herein obtained Seurat object were implemented for filtering, normalization, integration, clustering and differential expression analysis for different samples. Cells were filtered for genes expressing at least 100 cells. Normalization on cells was performed using ‘sctransform’ normalization method and integration of scRNA-seq data was performed on multiple samples using the top 3000 variable features. After integration, genes were used as input for executing principal component analysis (PCA) using function ‘RunPCA’ and genes were used for visualization 2D data in Uniform Manifold Approximation and Projection (UMAP) form using function ‘RunUMAP’. In Seurat, clustering was performed using the ‘FindClusters’ function adopting the resolution of 0.1 and the shared-nearest neighbor (SNN) graph was constructed using the first 30 PCA dimensions. In case of differential expression analysis, we used ‘FindAllMarkers’ with default parameters using negative binomial distribution (test.use = DESeq2) and minimum difference & minimum fraction between two groups with threshold of 0.1. Genes with log2FC above 1 and adjusted p-value less than 0.05 were considered as differentially expressed. To understand the enrichment of GO annotation from planaria, Swiss-prot blastx-id against the planarian annotation were aligned using the uniprot database. Furthermore, we also performed pseudo-time trajectory analysis and cell ordering through a pseudo-temporal continum using Monocle (v2) (Qiu et al., 2017) and Slingshot (v2) (Street et al., 2018). Gene ontology enrichment analysis for enriched genes was performed using EnrichR (E. Y. Chen et al., 2013).

### Construction of Gene Regulatory Networks

Gene regulatory networks (GRNs) were designed on the scRNA-seq data using R package hdWGCNA (v0.2.19) (Morabito et al., 2023) which incorporates the WGCNA (v1.72-1) to perform the co-expression network. Firstly, we constructed metacells to transcriptionally enable neighboring cells. hdWGCNA uses K-nearest neighbors (KNN) for building metacells. These obtained metacells were further processed for normalization by using function ‘NormalizeMetacells’ and after executing the function ‘TestSoftPowers’ appropriate co-expression networks were executed. We applied the AddModuleScore function in Seurat to compute the score for module related genes and obtain the expression pattern for our scRNA-seq datasets. In it module eigengenes were computed for gene expression profile in each sample alongside module connectivity were also calculated and average gene expression for each module were plotted in all clusters. Along with GRN analysis we also obtained Transcription factors (TF) from module-1 genes of co-expression network and TF from differentially expressed genes using ChEA3 (Keenan et al., 2019) with the GTEx library.

### RNA extraction and cDNA synthesis

To extract RNA, animals were kept in TRIzol (500ul) and stored at −80°C for overnight in 1.5ml Eppendorf tube. Using a pestle the tissue was homogenized and added 1/5^th^ volume of ice cold chloroform and mixed thouroughly using a pipette. The solution was kept on ice for 10 imn and centrifuged at 13,000 rpm for 30 mins at 4°C. the aquesous phase was transferred into a freshtube and pre-chilled isopropanol of equal volume is added and mixed. The solution was kept at −20°C for 1h/overnight. The solution was then centrifuged at13,000 rpm for 30 mins. The pelleted RNA was black in colourand washed with 70% ethanol (in nuclease-free water) twice. After washing, air dry the pellet and resuspend in nuclease free water. The RNA as stored at −80°C. 500 ng of RNA was used to prepare cDNA using TAKARA cDNA synthesis kit.

### dsRNA and Riboprobe preparation

Genes were amplified using gene-specific primers and cloned using the TA cloning kit. Primer sequences are mentioned in Tabel 5. XL10 *Escherichia coli* cells were used for transformation. By using blue-white colony screening to identify successful transformants in the presence of kanamycin resistance, positive clones were selected. Gene-specific primers with T7 promoter overhang were used to generate the template for dsRNA synthesis. Dig RNA labeling mix or Fluorescein RNA labeling mix was used to make riboprobes. Both dsRNA and riboprobe were generated using T7 polymerase-dependent in-vitro transcription. dsRNA was purified using the sodium acetate precipitation method and Riboprobe was purified using Megaclear transcription purification kit.

### RNAi

dsRNA was mixed with beef liver extract with a final concentration of 250 ng/μl. This feed was fed to one week-starved animals every 4^th^ day for 6 feeds. After the 6^th^ feed, animals were observed for one week for phenotype. Gene knockdowns (KD) that did not show any phenotype after 7 days were validated for KD using qRT-PCR. dsRNA of Green fluorescent protein (GFP) was used as the negative control.

### qRT-PCR

To perform qRT-PCR, the RNA was isolated using TRIzol after knockdown. The RNA was treated with DNase and then used 500ng of RNA to synthesise cDNA. PowerUP SYBRGreen master mix was used to perform Quantitative real-time PCR (qPCR). Run for every gene is done in triplicate using the QuantStudio 5 system. Refer to Table 6 for the primer sequences.

### Whole-mount *in situ* hybridization (ISH) and immunostaining (IHC)

Riboprobes were prepared using rNTPs labelled with either Digoxigenin or Fluorescein. Reverse primer with T7 overhang and the forward primer were used to amplify the gene. In it amplicon has been used to synthesize antisense Riboprobe. The used primers are listed in table.x. For both ISH and IHC, animals were killed using 5% N-Acetyl-L-Cysteine in pbstx for 5 min and fixed using 4% formaldehyde in pbstx (0.3% Triton-× 100, unless mentioned otherwise) for 20 mins. The fixed planaria were then dehydrated using methanol and stored at −20°C. Before staining, animals were rehydrated and bleached (5% deionized formamide, 0.5x SSX, 1.2% H2O2 in NFW) under bright light.

To perform in situ hybridisation, animals were permeabilised using proteinase-k and fixed using 4% formaldehyde. Riboprobe incubation was done in hybridization buffer for 16-18 hours at 56°C. The riboprobe concentrations used are 0.5 ng/μl for *smedwi-1,* 0.3 ng/μl for *ddx43,* and 0.5 ng/μl for *h1b.* Then the samples were washed with SSC, TNTx (10% 1M Tris-Cl pH: 7.5, XM NACL, 0.3% TritonX-100) and incubated with Anti-Digoxigenin POD (1:1000) or Anti-Fluorescein POD (1:1000) at 4°C overnight in blocking solution (5% RWBR, 5% Horse Serum in TNTx). The samples were washed with TNTX and developed using FITC/CY3/CY5 – tyramide signal Amplification (R. S. King & Newmark, 2013). For developing a second Riboprobe, the peroxidase enzyme on the first riboprobe antibody was inhibited using 0.215M sodium azide for 2 hours and further steps were continued form blocking. To perform immunostaining, the animals were washed with pbstx after bleaching and incubated in blocking solution (10% Horse serum in pbstx) for 2 hours. The primary antibody is then added with fresh blocking solution (1:100 for H3p) for 36-40 hours at 4°C. The samples are then washed with pbstx and incubated with secondary antibody (1:1000) in 10% Horse serum blocking solution. For BrdU staining, after the tyramide signal development the samples were treated with 2N HCL in 0.5% pbstx for 15 mins at 37°C. This is followed by primary (1:100) and secondary antibody(1:1000) incubations, as previously mentioned earlier in immunostaining.

### BrdU labeling strategy

Planaria were treated with a higher salt concentration (5x montjuic juice) for at least 2 days before BrdU feeding (Molinaro et al., 2021; Newmark & Sánchez Alvarado, 2000). To prepare BrdU stocks, 5mg of BrdU is dissolved in 100 μl of 50% DMSO in nuclease-free water and stored in −80. 20ul of this stock is then mixed with 80ul of beef liver and 100ul of 1% Ultra LowMelting Agarose. This Brdu feed (5mg/ml final concentration) was fed to planaria. Planaria were maintained in high salt concentration during the chase period. After the desired chase period, animals were are killed using 5% NAC and fixed with 4% formaldehyde as mentioned in the IHC section. Only the animals that didn’t developed any lesions during the chase period were fixed. BrdU staining was done in-addition to in situ hybridisation. After developing the probe, the animals were washed twice with pbstx. Samples were then incubated with 2N HCL in 0.5% pbstx at 37°C for 20 mins. The samples were then washed with pbstx and blocked using 10% Horse serum. Post blocking 1° antibody incubation was done for 36 hours at 4°C. Then 1° antibody was removed and samples are washed with pbstx multiple times. Secondary incubation was done in 10% HS at RT for 2 hours. Later samples were then washed and proceeded for DAPI staining.

### Irradiation

Cobalt based gamma irradiator (LDI-2000, Manufacturer: BRIT, INDIA) was used to irradiate planaria exposure time was calculated based on the dosage rate. Media is changed everyday and the animals are fixed using 4% Formaldehyde and stored in 100% methanol at −20°C.

### Inhibitor treatment

Erk inhibitor FR180204 and Notch inhibitor DAPT were both prepared as 50 mM stock solutions in 50% DMSO. Working concentrations of 25 µM (FR180204) and 35 µM (DAPT) were then made using Montjuïc water. Animals are pretreated for 4-hours with a freshly prepared inhibitor solution. Amputation was then performed just below the head. Following amputation, fresh inhibitor media was added to the animals, and they were fixed 6 hours later using 4% formaldehyde. Finally, samples were stored in 100% methanol at −20°C.

### Microscopy, Quantification and Software

Brightfield images were taken using Olympus SZX16 microscope. Confocal imaging was done on an FV3000 Olympus microscope with 40X 1.3 NA objective and 60X 1.42 NA objective. 3D reconstruction was done using FV3000 software from Olympus. Quantification of immunostaining and FISH stainings were done using Fiji software (V 1.54p). Statistical significance and Graphs were generated using Graphpad PRISM software. Models were made using Biorender.

### Bulk RNA Sequencing and Analysis

Animals were fed with *ddx43* dsRNA for 6 feeds and RNA was isolated after 7 days of the last feed. The RNA was isolated using TriZOL and coprecipitated using glycoblue before submitting for sequencing. Samples were sequenced in triplicate. mRNA was isolated using NEBNext® Poly(A) mRNA Magnetic Isolation. Library preparation was done using NEBNext® Ultra™ II Directional RNA Library Prep Kit and sequenced on Illumina Nova seq 6000. Raw RNA-seq reads were subjected to quality control and trimming using fastp [v0.20.1] (S. Chen et al., 2018) followed by aggregation of the QC report data in MultiQC [v1.9] (Ewels et al., 2016). STAR [v2.7.9a] (Dobin et al., 2013) was used for mapping using *S. mediterranea (*S2F2) as a reference genome. Mapped reads were processed to obtain raw counts for gene expression estimation using featureCounts (Liao et al., 2014) from the subread package [v1.5.2]. Differential gene expression was analyzed for differential expression using DESeq2 [v1.30.1] (Love et al., 2014) in the R environment [v4.0.3]. In order to determine upregulated and downregulated genes for *ddx43* knockdown and GFP control, differentially expressed genes with an absolute value of log2foldchange greater than 1 and false discover rate less than 0.05 were filtered using general linearized model. Heatmap were created using default parameters by obtaining z-scores from normalized read counts generated by DESeq2.

## Acknowledgements

We would like to thank Dr Tina Mukherjee and Dr Kai Lei (Wuhan University) for their discussion and valuable feedback on our manuscript. We acknowledge the central imaging and flow cytometry facility (CIFF), NGS facility and Irradiation facility at the BLiSc Bio-Cluster for their constant technical support. We also thank Johan from Colin lab for Erk Inhibitor and Raj lab (NCBS) for DAPT.

## Competing Interests

The Authors declare no competing or financial interests.

## Author contributions

N.K.J., M.M.H., A.A., and D.P. contributed to the conceptualization of the study and manuscript preparation. N.K.J. performed the majority of the experiments and supervised all experimental work. Confocal imaging was carried out by N.K.J. Single-cell isolation and sequencing were performed by M.M.H., with assistance from P.V. All bioinformatic analyses were conducted by A.A. BrdU experiments were performed by M.B. Radiation experiments were conducted by A.M. and V.K.D. M.B., A.M., and S.P. assisted with in situ hybridization and immunostaining. D.P. acquired funding and supervised the project.

## Resources

Single cell X1 data used in this paper are taken from NCBI GEO: GSE107873

## Funding

N.K.J and A.M would like to thank Department of Biotechnology (DBT), India and Institute for Stem Cell Science and Regenerative Medicine (inStem) for PhD fellowship. This work was supported by inStem core grant, Swarnajayanti fellowship and Frontiers grant.

## Supplementary Figure legends

**Figure S1:**
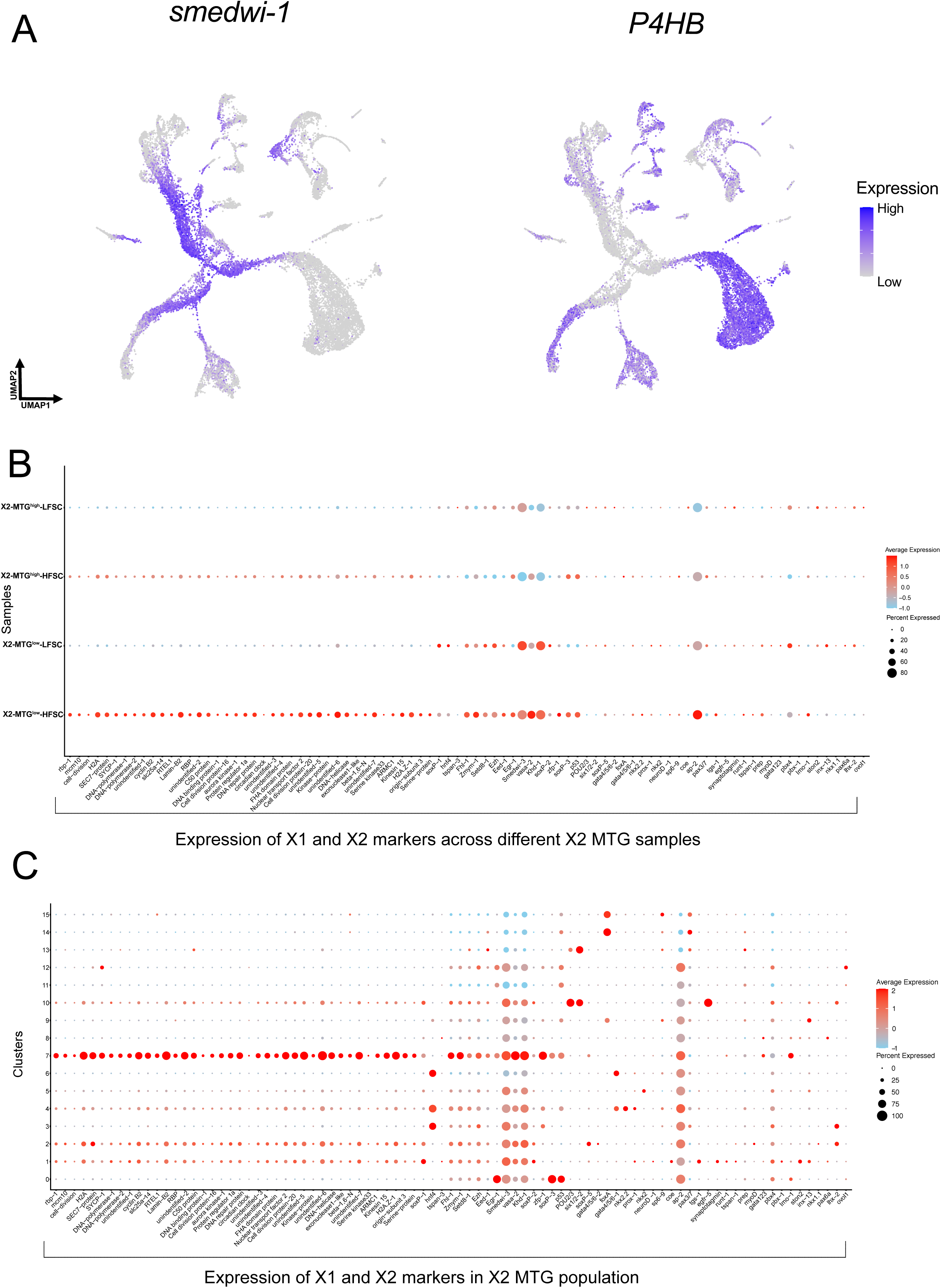
scRNA-seq entraps X1 and X2 markers expression over MTG population. (A) UMAP expression representation for neoblast marker as *smedwi-1* and *P4HB* as post-mitotic marker (B) Dot plot illustration for previously published X1 and X2 population markers obtained from Raz et al., (2021) across different X2 MTG samples. In it *mcm-10, ap-2, cyclin B2, lamin B2, vasa-2 and sycp-1* have shown to be expressed in pluripotent (X2 MTG^low^ HFSC) sample (C) Dot plot identification of published X1 and X2 population markers obtained along clusters from X2 MTG population. In it cluster 7 seems to represent higher expression for markers like *mcm-10, ap-2, cyclin B2, lamin B2, vasa-2 and sycp-1*.

**Figure S2:**
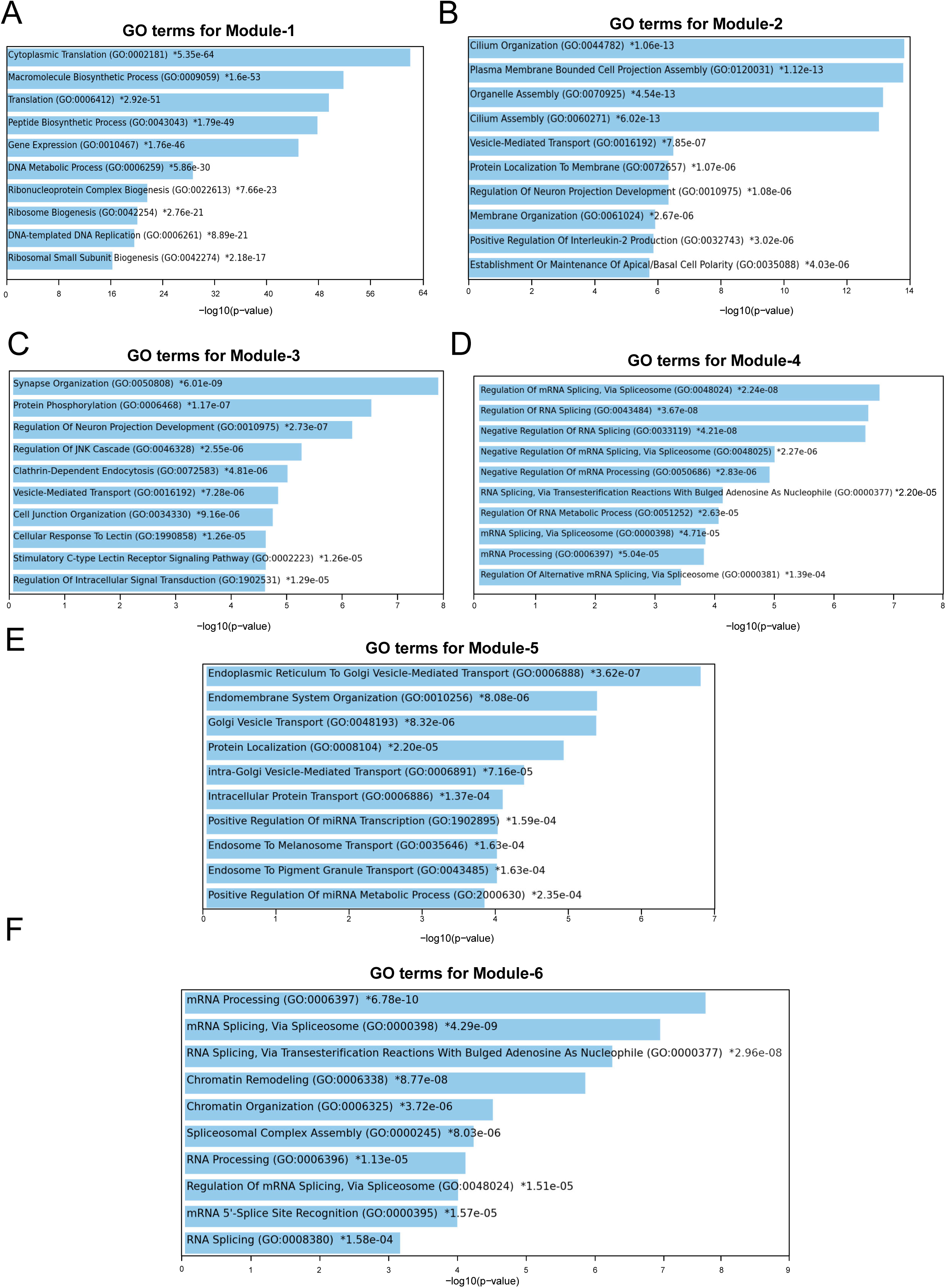
Enrichment analysis for identified GRN modules. (A) Bar plot representing top 10 gene ontology terms for the genes obtained from GRN analysis for each specific module. In module-1, Cytoplasmic Translation and Ribosome biogenesis appear to be highlighted terms. While star(*) mentioned in respective terms constitute adjusted p-value less than 0.05 (B) Bar plot identifying obtained top 10 gene ontology terms for the genes obtained in module-2. In it, Cilium assembly, membrane organization and maintenance of basal cell polarity appeared to be enriched terms from it (C) Bar plot for characterization of module-3 revealed terms like Synapse organization, Neuron projection, Cell junction organization and Intracellular signal transduction for its set of genes (D) Bar plot outlining enriched terms from module-4 genes such as RNA splicing, regulation of RNA metabolic process and mRNA processing (E) Bar plot illustrating enriched terms from module-5 genes such as Endoplasmic reticulum to golgi vesicle mediated transport, Protein localization and Intracellular protein transport (F) Module-6 genes exhibiting GO terms like Chromatin Remodeling, Chromatin Organization and RNA splicing

**Figure S3(1):**
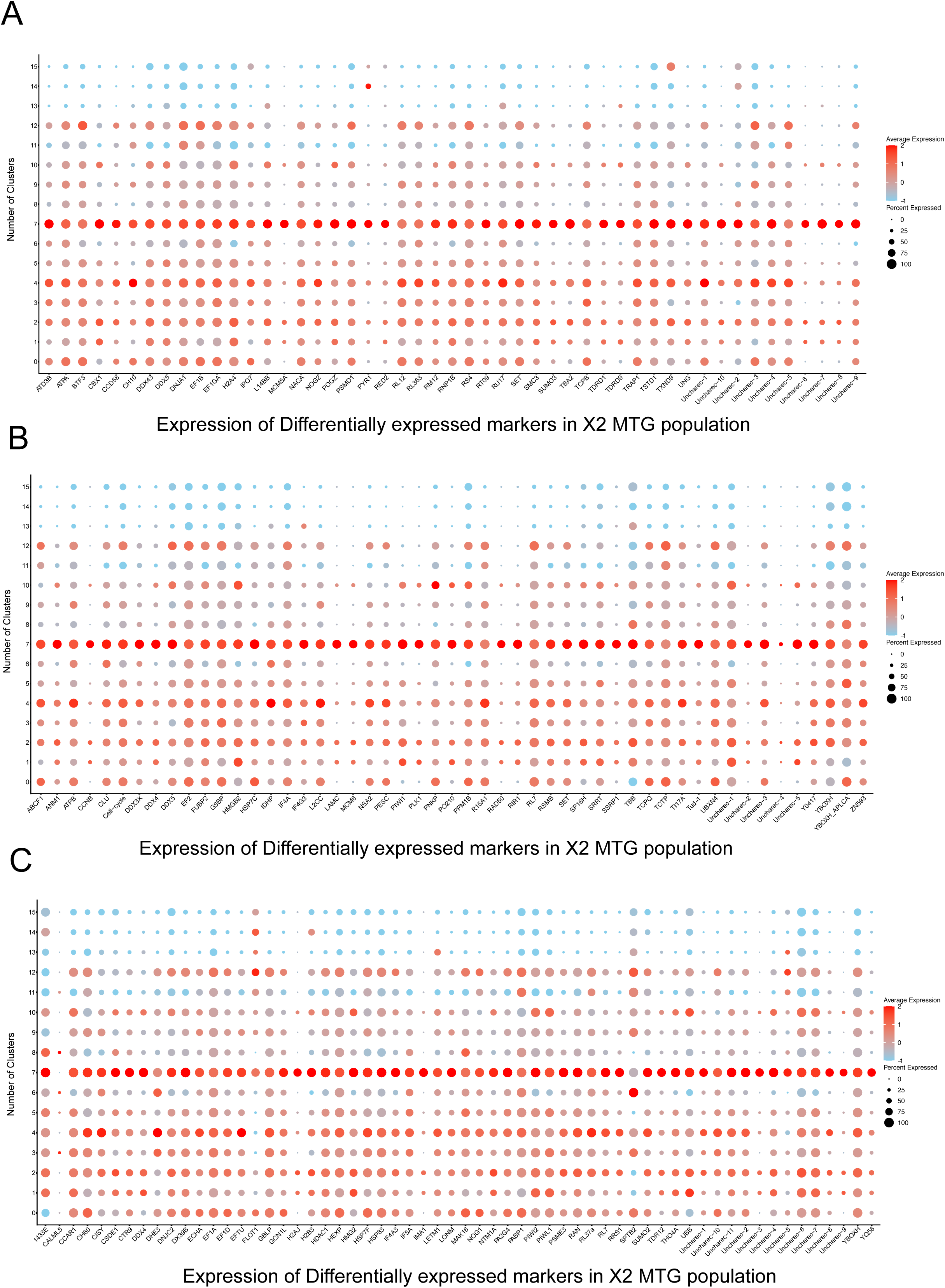
Expression pattern of genes across X2 MTG samples derived clusters. (A) Dot plot identifying the expression from first set differentially expressed gene across X2 MTG population. Differentially expressed genes (n=209) were distributed into different sets for plotting across different population. (B,C) Differentially expressed genes from second and third set across X2 MTG population

**Figure S3(2):**
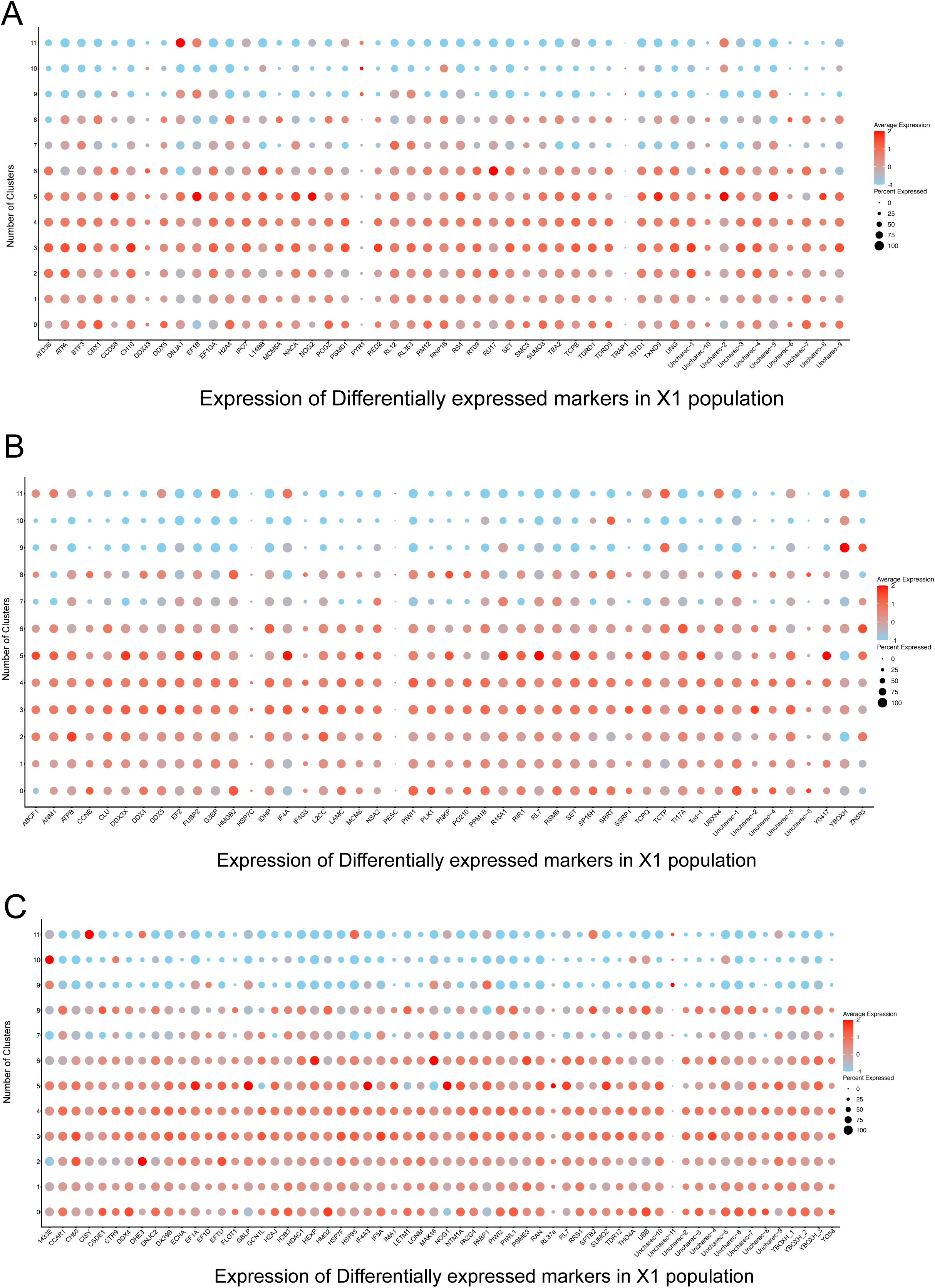
Expression pattern of genes across X1 population derived clusters. (A) Dot plot depicting the expression from first set of differentially expressed genes across X1 population (B,C) Differentially expressed genes in the form second and third set across X1 population

**Figure S3(3):**
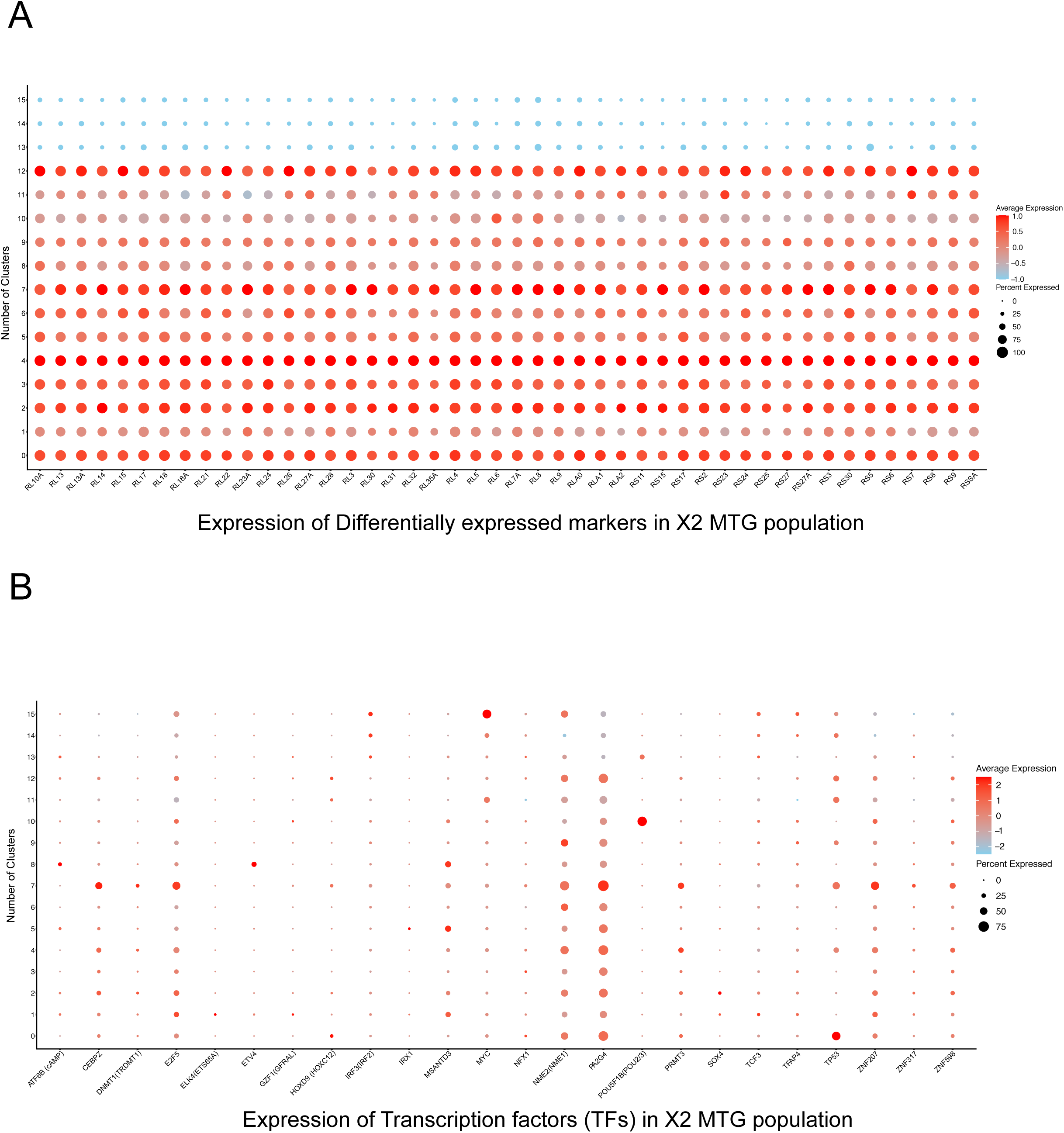
Expression pattern of genes across X2 MTG samples derived clusters. (A) Dot plot for fourth set of differentially expressed genes across cell clusters of X2 MTG population (B) Transcription factors (TF) obtained from GRN and Differential expression analysis were considered for expression quantification over X2 MTG clusters

**Figure S3(4):**
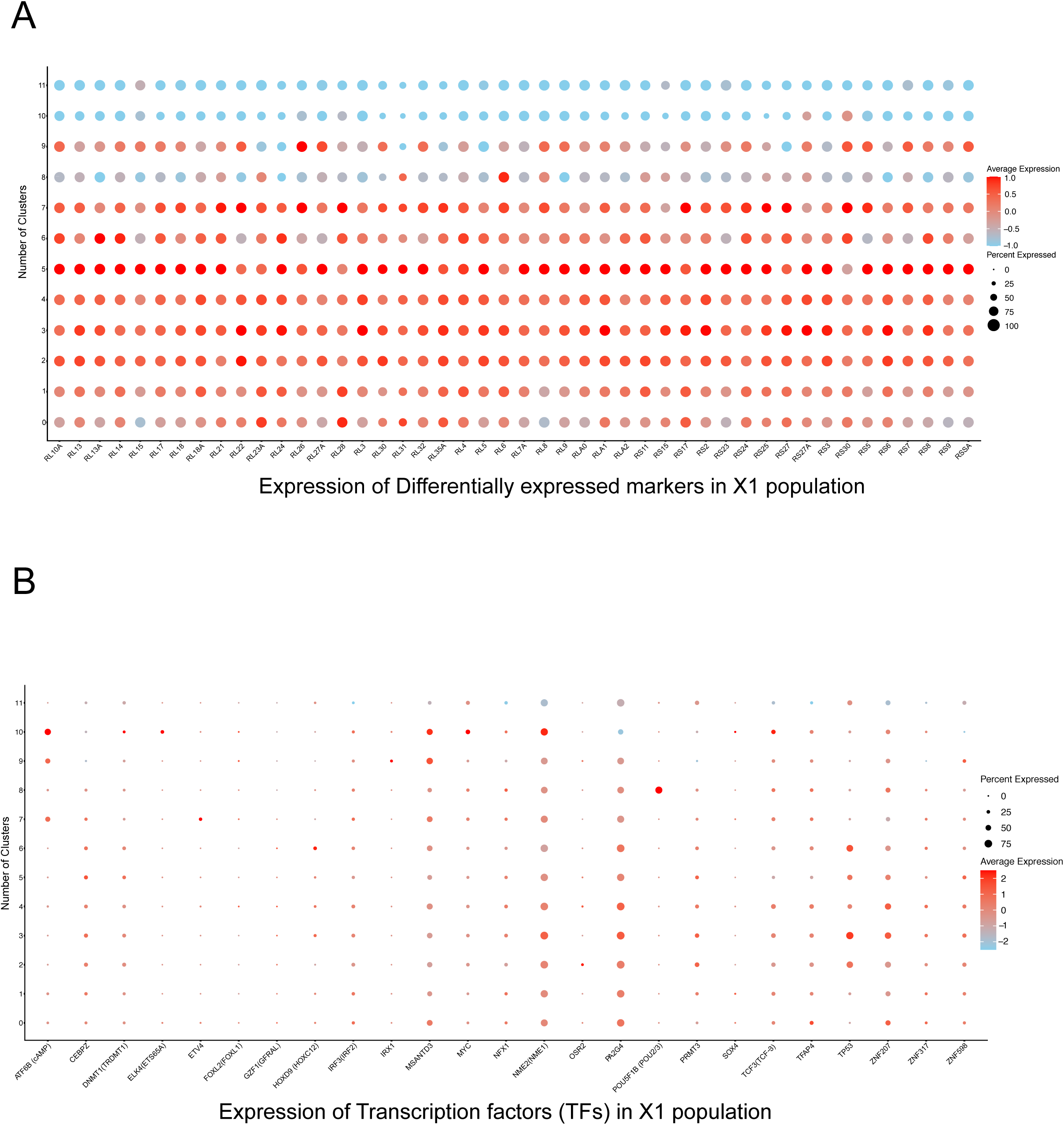
Expression pattern of genes across X1 population derived clusters. (A) Fourth set of differentially expressed genes were represented using dot plot (B) Similar set of TF were used for expression computation across cell clusters of X1 population

**Figure S3(5):**
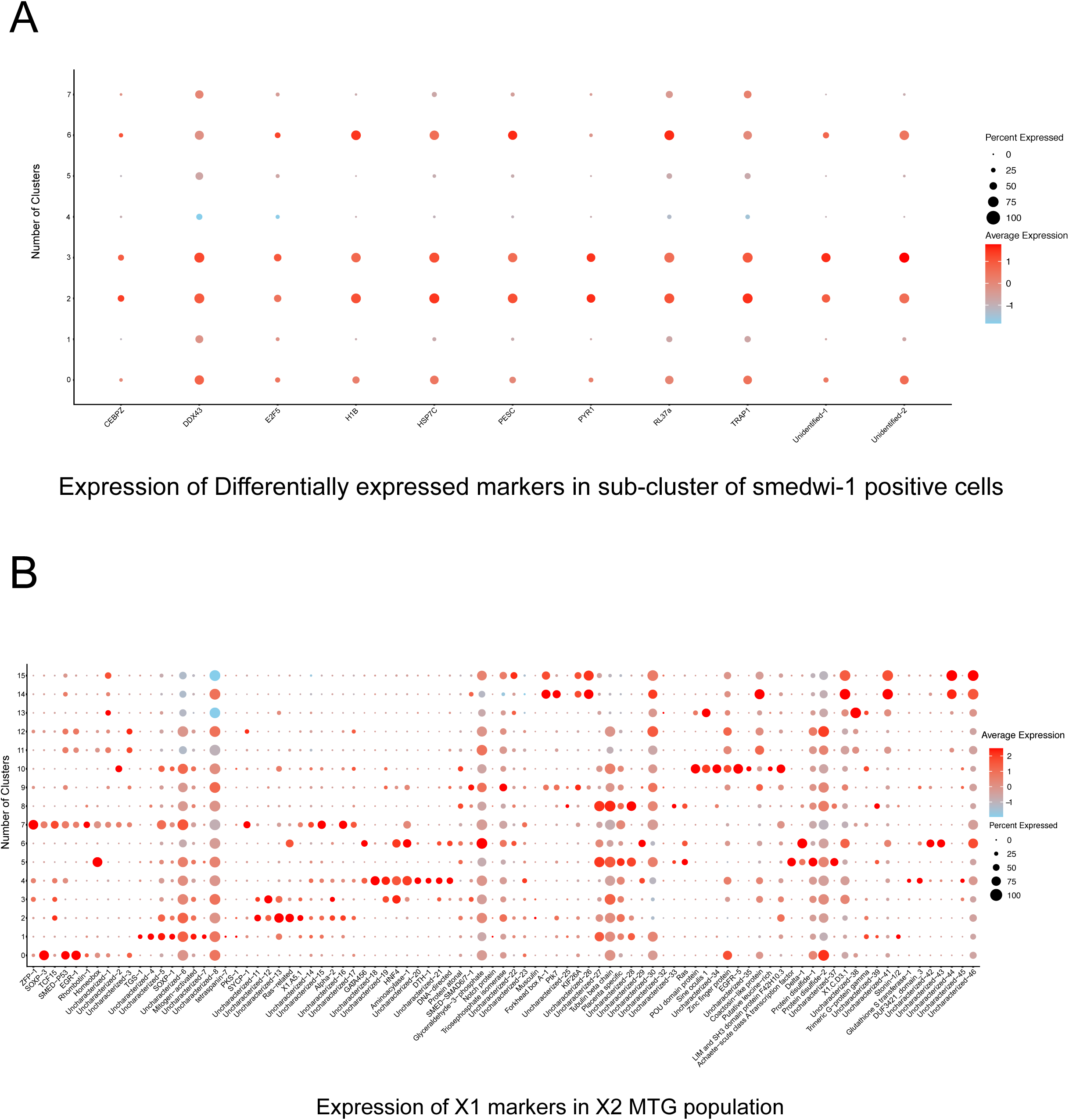
Expression outline of X2 gained markers in sub-clustering of neoblast cluster. (A) Dot plot describing the eleven enriched markers over sub clustering of neoblast cluster. In it, Cluster 0,2,3 and 6 portrays enriched cluster for obtained markers (B) Dot plot depicting the expression levels of previously published X1 population markers over scRNA-seq data of X2 MTG population. From it markers namely *zfp-1*, *sycp-1, rhombotin-1* seems to be enriched in neoblast cluster

**Figure S4:**
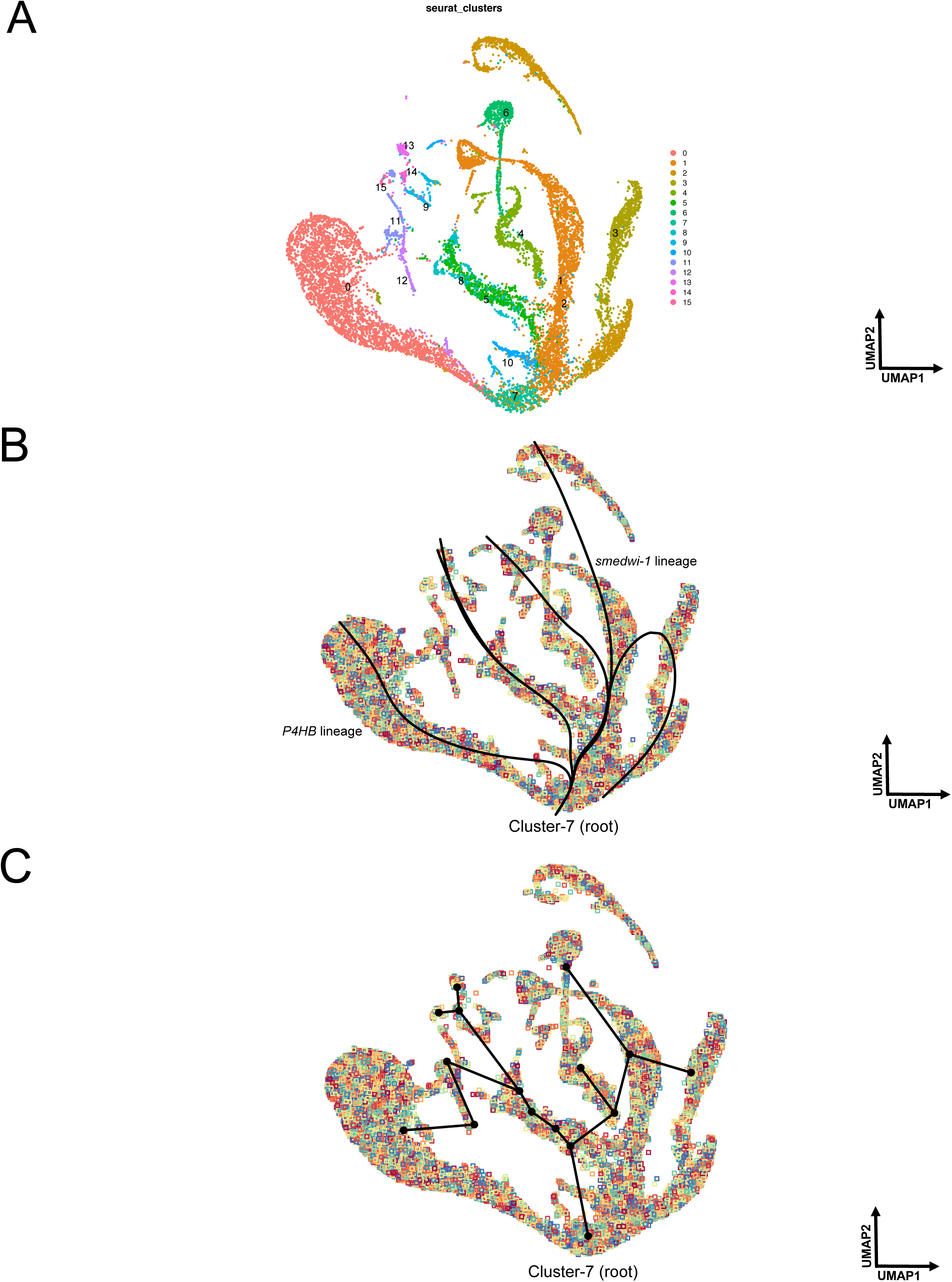
scRNA-seq identifies trajectory inference from intergal cells. (A) UMAP plot illustrating the trajectory inference taking cluster 7 (neoblast cluster) as root for distribution of total cells across 16 clusters using slingshot (B) UMAP plot identifying the two different lineages across our total cells from root cluster. *smedwi-1* lineage shown on the left and *P4HB* lineage shown on the right. (C) UMAP plot describing the smooth trajectory using the principal curves from the obtained lineages by taking neoblast cluster as root

**Figure S5:**
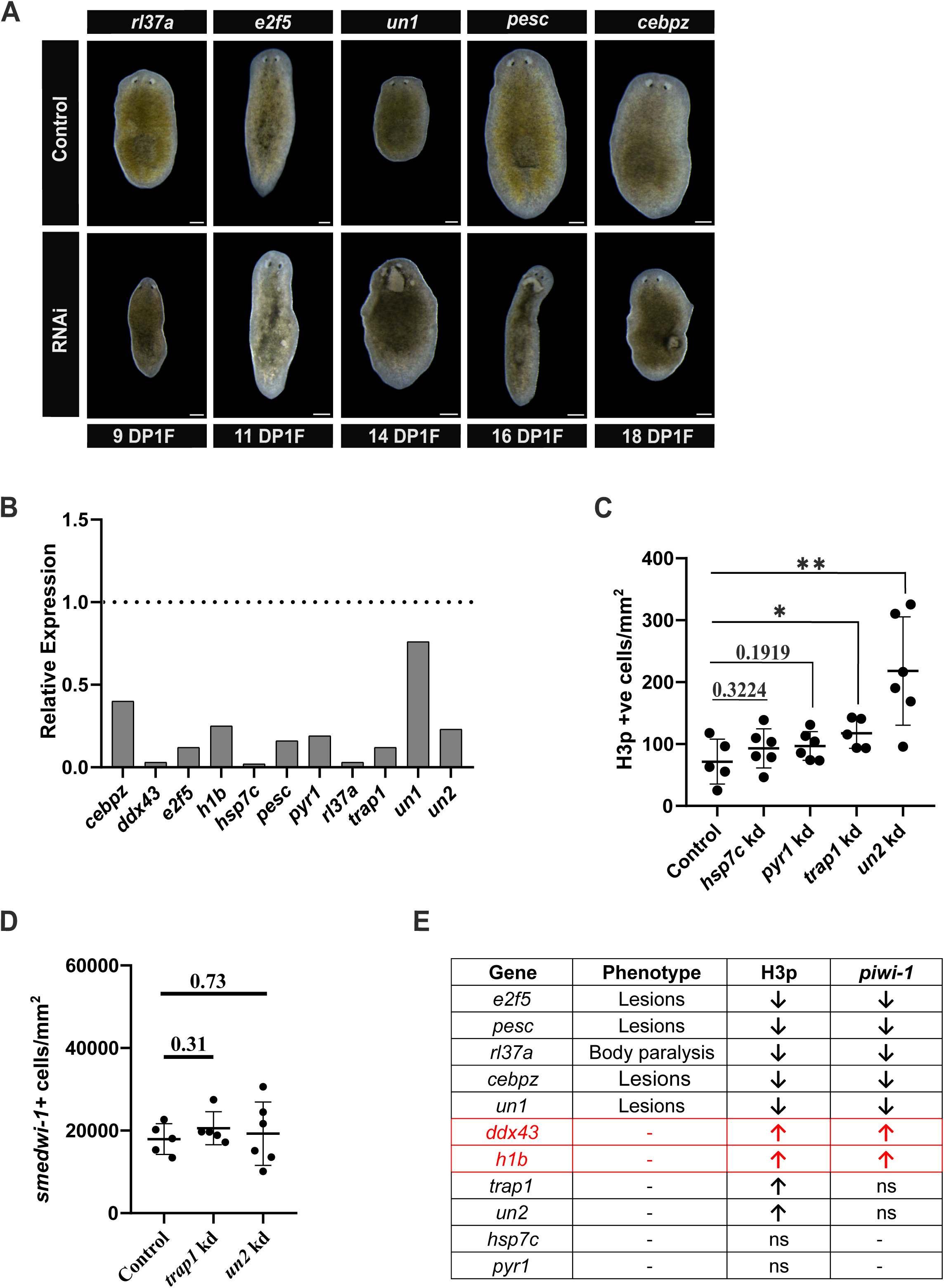
Ectopic phenotype observation for enriched markers. (A) Darkfield images of *rl37a, e2f5, un1, pesc*, and *cebpz* knockdowns at 9, 11, 14, 16, and 18 days post first feeding (DP1F). (B) qPCR analysis showing relative gene expression for 11 knockdowns. (C) Quantification for H3p^+^ cells in *hsp7c, pyr1, trap1*, and *un2* knockdown animals. p value: *<0.05; ** <0.01 (D) Quantification for *smedwi*-1^+^ cells in *trap1* and *un2* knockdown animals. (E) Table summarizing obtained phenotypes, H3p^+^ and *smedwi-1*^+^ cell quantification description for 11 genes.

**Figure S6.**
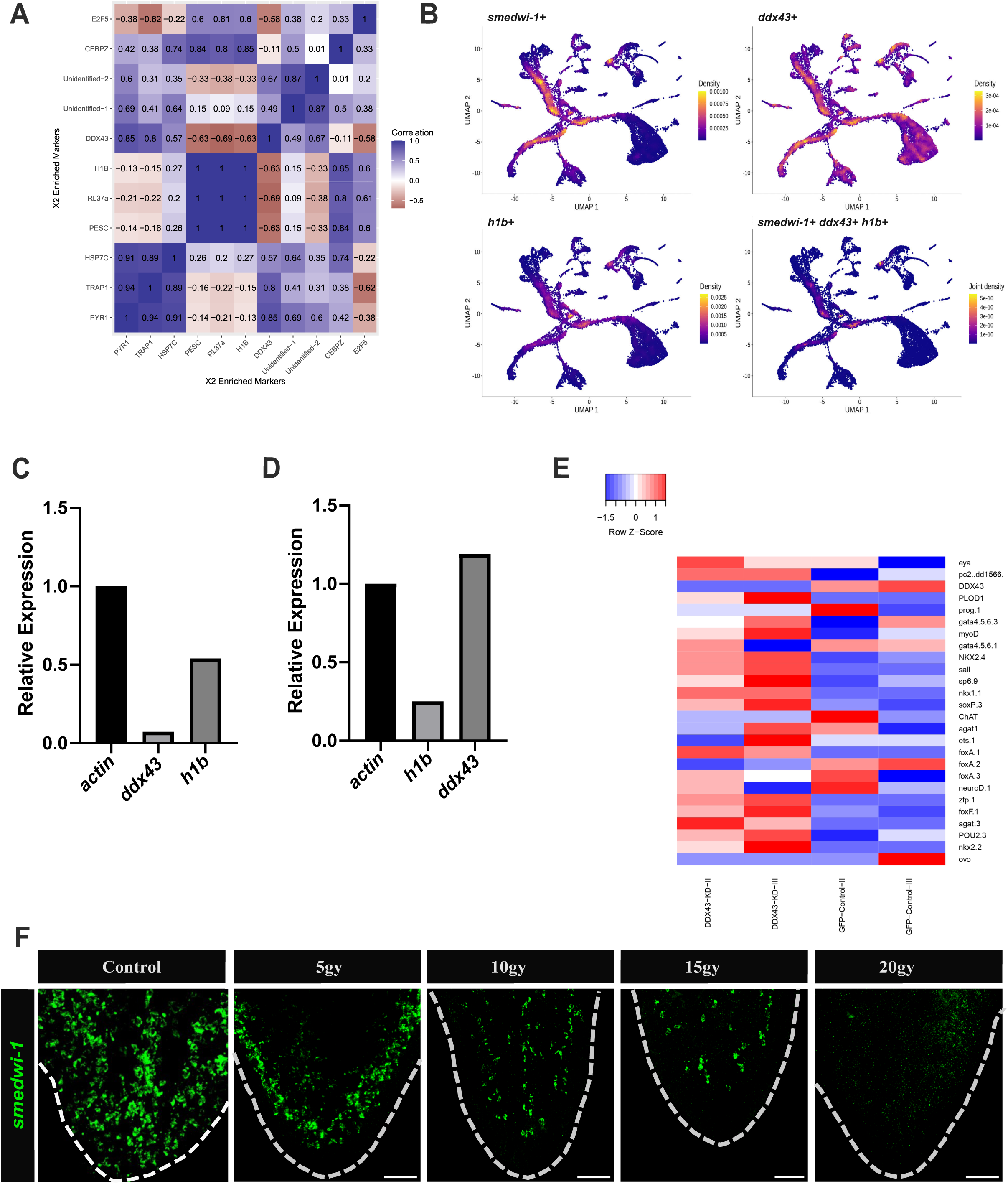
Bulk RNA-seq and scRNA-seq captures interaction for X2 enriched markers. (A) Correlation coefficient heatmap between enriched markers featured in pluripotent sample of X2 MTG^low^ HFSC obtained from differential expression with other progenitors sample in X2 MTG population. In it blue color represents the positive correlation while the brown color represents the negative correlation. (B) UMAP plot illustrating the positive cells in form of higher density for respective markers like *smedwi-1, ddx43* and *h1b*. All together it represents triple positive cells including markers *smedwi-1, ddx43* and *h1b* for quantification in form of joint density. (C) Relative expression of *ddx43* and *h1b* in *ddx43* knockdown animals. (D) Relative expression of *h1b* and *ddx43* in *h1b* knockdown animals. (E) Heatmap showing expression pattern of previously published differentiation markers from H. O. King et al., (2024) and Raz et al., (2021) across different replicates for GFP control and *ddx43* RNAi sample using normalized read counts (z-score). In it, red color depicts the higher expression and blue color depicts the lower expression. (F) RNA-FISH showing *smedwi-1^+^* cells in wild-type and irradiated animals exposed to 5, 10, 15, and 20 Gy respectively.

**Figure S7:**
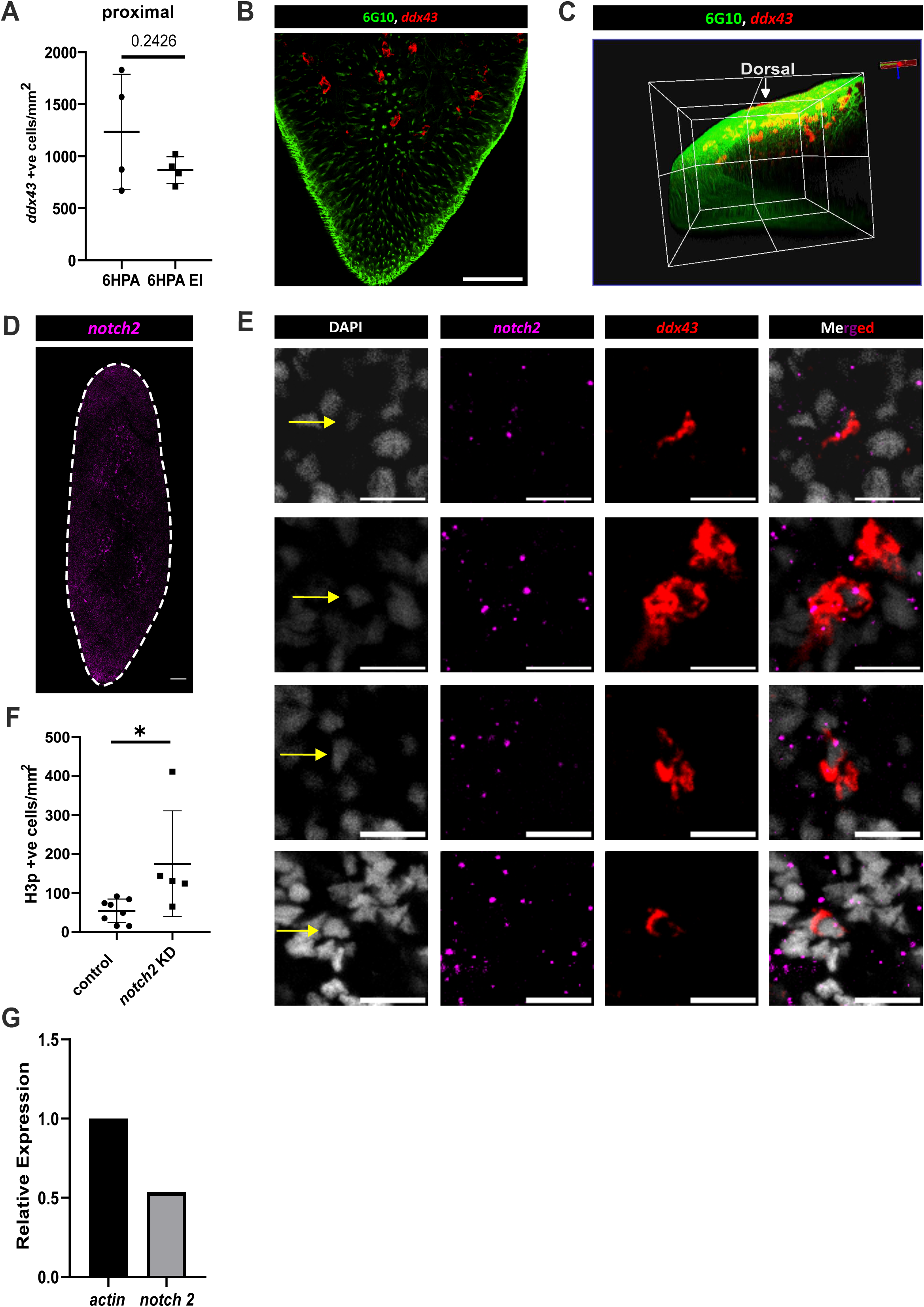
Notch activity required for ddx43 cell maintenance. (A) Quantification of *ddx43*^+^ cells in the proximal region at 6 hpa in untreated and Erk-inhibited animals. Each condition comprising four animals. (B,C) RNA-FISH for *ddx43* and immunostaining with 6G10 (muscle marker) shows *ddx43*^+^ cells closely associated with sub epidermal muscles on the dorsal side. Scale bars: 100 µm. (D) RNA-FISH showing *notch2* expression in wild-type animals. Scale bars: 100 µm. (E) RNA-FISH for *ddx43* and *notch2* in wild-type animals. *notch2* expression was observed in *ddx43^+^* cells. Scale bars: 10 µm. (F) Quantification indicating H3p^+^ cells in *notch2* knockdown animals. pvalue: *<0.05.

